# Olfactory learning primes the heat shock transcription factor HSF-1 to enhance the expression of molecular chaperone genes in *C. elegans*

**DOI:** 10.1101/152736

**Authors:** Felicia K. Ooi, Veena Prahlad

## Abstract

Learning, a process by which animals modify their behavior as a result of experience, allows organisms to synthesize information from their surroundings to acquire resources and predict danger. Here we show that prior encounter with the odor of pathogenic bacteria prepares *Caenorhabditis elegans* to survive actual exposure to the pathogen by increasing HSF-1-dependent expression of genes encoding molecular chaperones. Learning-mediated enhancement of chaperone gene expression requires serotonin. Serotonin primes HSF-1 to enhance the expression of molecular chaperone genes by promoting its localization to RNA polymerase II–enriched nuclear loci, even prior to transcription. HSF-1-dependent chaperone gene expression ensues, however, only if and when animals encounter the pathogen. Thus, learning equips *C. elegans* to better survive environmental dangers by pre-emptively and specifically initiating transcriptional mechanisms throughout the whole organism. These studies provide one plausible basis for the protective role of environmental enrichment in disease.

## Introduction

The ability to accurately predict danger and implement appropriate protective responses is critical for survival. Many animals possess neuronal circuits to detect unfavorable conditions and implement an avoidance response. In addition, all cells possess conserved mechanisms to repair and protect their macromolecules from damage that occurs under adverse conditions. One such mechanism present in all cells to protect against protein damage is the Heat Shock Response (HSR) (1-4). The HSR is mediated by the transcription factor Heat Shock Factor 1 (HSF1), which, in response to a variety of stressors, increases the expression of cytoprotective molecular chaperones or so-called heat shock proteins (HSPs) to maintain protein stability and help degrade proteins that misfold and aggregate under stressful conditions (1-4). HSF1 activity is essential for all organisms to adapt to changing environments. The HSR and HSF1 can be activated autonomously by single cells in response to proteotoxic stressors (1-3). However, within a metazoan such as the nematode *Caenorhabditis elegans*, HSF1 and the cellular response to protein damage are not autonomously controlled by individual cells, but instead are under the regulation of the animals’ nervous system (5-11). The biological role for this regulation is unclear. We discovered that one mechanism by which *C. elegans* HSF1 (HSF-1) is regulated is through the neurosensory release of the bioamine serotonin (5-hydroxytryptamine, 5-HT (7)). In vertebrates and invertebrates, serotonergic systems play a central role in neurophysiological processes underlying learning and memory, allowing animals to learn about threats in their environment and form memories that can be later recalled to modify behavior (12-23). Therefore, we asked whether control by the serotonergic-based learning circuitry allowed *C. elegans* to modulate HSF-1 activity in response to prior experience, so as to better combat threats in its environment.

Here we show that in *C. elegans*, olfactory experience of specific odorants released by the toxic bacteria *Pseudomonas aeruginosa* PA14 primes HSF-1-dependent transcription of cytoprotective *hsp* genes, such that the expression of these genes is enhanced if and when animals subsequently encounter the pathogen. This priming requires 5-HT and occurs through a novel mechanism, whereby HSF-1 is pre-emptively mobilized to the vicinity of RNA polymerase II (pol II) in nuclei throughout the animal, in preparation for active transcription. Animals that cannot synthesize 5-HT are deficient in re-localizing HSF-1 in response to olfactory stimuli, and do not show this learned enhancement of *hsp* expression. Olfactory priming of HSF-1 is protective, allowing animals that had previously experienced the smell of *P. aeruginosa* to better respond to a subsequent exposure to the pathogen. Thus neuronal control over the HSF-1-mediated defense mechanism of cells allows learning and memory to elicit anticipatory changes in the stress-responsiveness of cells, facilitating survival. X

## Results

### Olfactory exposure to odorants made by the toxic bacteria Pseudomonas aeruginosa PA14 accelerates the avoidance response of C. elegans to the pathogen

To test whether animals can ‘learn’ to activate HSF-1 based on prior experience, we exploited previous findings that *C. elegans* have an innate aversion to specific pathogens, and display experience-dependent plasticity to avoid ingesting pathogenic bacteria such as *Pseudomonas aeruginosa* PA14 (15, 24). Thus, although animals are typically attracted to novel bacteria upon initial encounter, be it pathogenic bacteria such as PA14 or non-pathogenic *E. coli* strains (25, 26), animals previously exposed to a lawn of pathogenic PA14 will avoid PA14 lawns upon subsequent exposure. This learned avoidance behavior requires the olfactory nervous system and 5-HT (15, 24, 27, 28). We used this information to set up a paradigm whereby we could train animals to avoid PA14 using odor alone, circumventing any physical damage that could be inflicted by actual exposure (Fig. S1A). We then asked if olfactory training on the odorant of this toxic bacteria could enhance the transcriptional activity of HSF-1 and promote survival if animals were to subsequently encounter the pathogen. Animals were trained by exposing them to the odor of PA14 cultures for 30 minutes. Controls were mock-trained by exposure to the odor of the standard *E. coli* OP50 strain on which animals are typically raised. To assess whether olfactory pre-exposure was sufficient to elicit learned avoidance behavior, trained and mock-trained animals were then given a choice between PA14 lawns and OP50 lawns. Behavioral preference was quantified by calculating a choice index (CI) for PA14, wherein a CI of 1.0 indicates maximal preference and a CI of −1.0 indicates maximal aversion (Fig. S1A). Due to the variability inherent to behavioral assays, all avoidance assays were conducted, and are represented, as pairwise comparisons between control and experimental populations of *C. elegans* evaluated in parallel. As previously reported (15), when faced with a choice between OP50 or PA14, naïve animals initially preferred the novel bacteria and accumulated within the first 5 minutes on PA14 (Fig. S1B). However, after 45 minutes *C. elegans* began to avoid PA14 and by 4 hours, 80% of the animals have left PA14 and moved to the OP50 lawn (Fig. S1B). Animals exposed to OP50 odor (mock-trained, control animals) behaved like naïve animals and also initially accumulated on PA14 and then began to leave the lawn by 1 hour (Fig. 1A). In contrast, animals exposed to the odor of PA14 avoided the PA14 lawn significantly earlier and left within the first 5 minutes (Fig. 1A). The avoidance of PA14 following pre-exposure to PA14 odorants appeared to reflect an innate response of the animals to PA14. It was also not a simple consequence of adaptation to the smell. This was inferred from the behavior of animals exposed for similar durations to the odor of another novel, but non-pathogenic bacteria, HT115. In this case animals did not avoid HT115 when given a choice between HT115 and OP50 but remained on HT115 throughout the analysis (Fig. S1C). This enhanced avoidance response was also specific to the pathogen in that animals responded to the pathogen whose odorants they had previously experienced. Pre-exposure of animals to the odor of PA14 did not trigger avoidance of another known *C. elegans* pathogen, *Serratia marcescens* strain DB11: animals pre-exposed to PA14 or OP50 odorants behaved like naïve animals and remained on DB11 throughout the analysis (Fig. S1D). These data taken together point to the existence of sophisticated mechanisms by which *C. elegans* discriminate between bacteria in their environment, and show that prior exposure to odorants generated by a pathogen such as *P. aeruginosa* can induce *C. elegans* to accelerate their avoidance of that specific pathogen upon subsequent encounter.

**Fig. 1.**
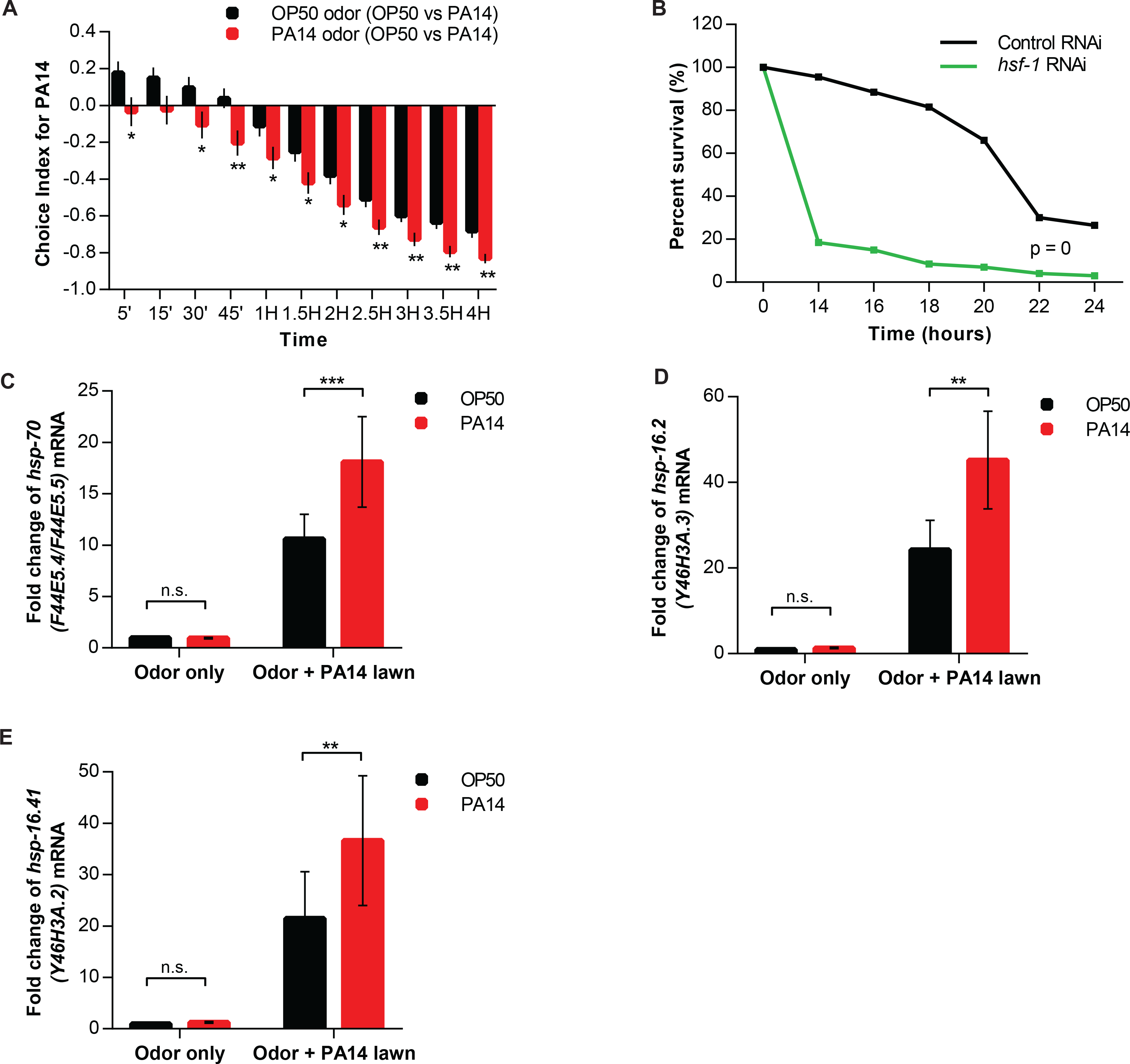
sNucDrop-Seq: a massively parallel single-nucleus RNA-Seq method. **(A)** Choice index for PA14 of wild-type animals pre-exposed to the odor of either OP50 or PA14. Preference was recorded at the times indicated on the x-axis. N = 16-17 experiments of 30 animals per condition. Student’s paired t-test. *p<0.05, **p<0.01. Legends: pre-exposure conditions (choice). **(B)** Survival on PA14 of ‘control’ animals and animals where *hsf-1* was knocked-down by RNAi. RNAi was conducted using standard methods for feeding RNAi (see Materials and Methods). N = 3 experiments of 50 animals per condition. Log-rank test. p=0. Also see Table S1. **(C–E)** *hsp-70* (F44E5.4**/**F44E5.5), *hsp-16.2* (Y46H3A.3), and *hsp-16.41* (Y46H3A.2) mRNA abundance measured by qRT-PCR upon exposing animals that had been trained on OP50 or PA14 odor to a lawn of PA14. Values were normalized to wild-type animals pre-exposed to OP50 odor. N = 38 (C), 12 (D), and 10 (E) experiments of 30 animals per condition. Pairwise mean comparison from linear mixed model analysis. **p<0.01, ***p<0.001. See Materials and Methods and Table S2 for complete details. (A, C, D and E) Data represent means ± S.E.M.

### The HSF-1-dependent expression of heat shock protein genes is enhanced by prior olfactory exposure to PA14 odorants

Exposure to PA14 is known to be toxic, causing increased protein damage (29, 30) and ultimately, death (31, 32). Consistent with this, survival on PA14 was HSF-1-dependent as the knock-down of *hsf-1* by RNA interference using standard methods for feeding double stranded RNA to *C. elegans*, accelerated death upon PA14 exposure (Fig. 1B; Table S1, also see Materials and Methods and Fig. S4A for confirmation of RNAi induced knockdown). To assess whether training by PA14 odorants modulated the HSF-1 transcriptional response, we placed animals exposed to OP50 odorants (controls) or PA14 odorants on PA14 lawns that covered the surface area that the animals explored, so animals could not implement their avoidance response. Under these conditions, HSF-1 was indeed activated: all animals placed on PA14 lawns for only 10 minutes increased HSF-1-dependent expression of the inducible *hsp70* F44E5.4/F44E5.5 genes, and the small heat shock proteins *hsp-16.2* and *hsp-16.41* as measured using qRT-PCR (Figs. 1C-E and Fig. S1E). Pre-exposure to the odor of PA14, however, enhanced this HSF-1-dependent transcriptional response (Figs. 1C-E; Table S2): the amounts of all three chaperone mRNAs were approximately two-fold higher in animals that were first pre-exposed to the odor of PA14, compared to control animals pre-exposed to the smell of OP50 (Figs. 1C-E; Table S2). This suggested that HSF-1–mediated gene expression could be enhanced by prior experience of signals that were predictive of danger. Of note, pre-exposure to the odor of PA14 did not, in itself, induce chaperone expression and animals exposed to the odor of PA14 had low basal chaperone expression similar to control animals (Figs. 1C-E; Table S2).

*P. aeruginosa* secretes several molecules that alter the behavior of other organisms. The “grape-like” odorant 2-aminoacetophenone (2AA) is one such compound synthesized relatively early in the growth cycle, and is enriched when *P. aeruginosa* infects animal tissue, such as wounds of human burn victims or the lungs of patients with cystic fibrosis (33, 34). 2AA is responsible for the attractive behavior of *Drosophila melanogaster* towards the pathogen (33, 35) as well as the aversive behavior of vertebrate species such as birds and mice from *Pseudomonas* (36, 37). We tested whether this compound was, at least in part, responsible for the learned enhanced aversion of *C. elegans* to PA14. Pre-exposure to 2AA mimicked the results observed in our choice assay although 2AA did not in itself elicit an avoidance response (Fig. S2A): animals that were pre-exposed to the smell of 2AA avoided PA14 lawns by 15 minutes compared to mock-trained, control animals exposed to the ‘odor’ of water who only began to avoid PA14 lawns by 45 minutes (Fig. 2A). Pre-exposure to 2AA odorant also enhanced the expression of *hsp70* mRNA when animals were subsequently exposed to PA14 lawns, although in itself, 2AA did not induce *hsp70* mRNA (Fig. 2B; Table S2). Moreover, the avoidance of 2AA did not appear to be due to its potential toxicity and prolonged 2AA exposure had no effect on the lifespan of animals, be it administered as an odor alone (Fig. 2C; Table S3), or mixed into OP50 for direct contact or ingestion (Fig. 2D; Table S3). The enhancement of PA14-avoidance behavior by 2AA also appeared to be fairly specific: pre-exposure to another volatile semiochemical secreted by *Pseudomonas*, N-3-oxododecanoyl homoserine lactone (3OC12-HSL(38)), did not affect subsequent avoidance behavior to PA14 lawns or enhance *hsp* gene expression (Figs. S2B, S2C; Table S2). Consistent with a role in signaling a potential threat, pre-exposure to 2AA appeared to facilitate a mechanism by which animals ‘decided’ to activate HSF-1 only if they subsequently encountered PA14, but not in its absence. This was seen when *C. elegans* that were pre-exposed to 2AA odorant encountered an OP50 lawn instead of a PA14 lawn: under these conditions, they did not activate HSF-1-dependent *hsp* gene expression (Fig. 2B; Table S2). However, if animals did encounter PA14, pre-exposure to PA14 odorants conferred a consistent and significant survival advantage: 63% of the animals pre-exposed to PA14 odor survived after 18 hours of PA14 exposure, compared to 46% of control, water-exposed animals (Fig. 2E; Table S4). The protection conferred by pre-exposure to PA14 was also stressor-specific, enhancing survival on PA14 but not upon prolonged heat stress (Fig. S2D). These data taken together suggest that the prior experience of PA14 odor, mimicked in large part by the odorant 2AA, was enhancing the organism’s ability to survive, not only by hastening the avoidance behavior of the animal from the pathogen, but also by enhancing the expression of cytoprotective HSF-1 transcriptional targets upon actual encounter with the pathogen. 2AA was in itself not toxic, nor aversive, but instead appeared to convey specific information regarding the bacterial environment of *C. elegans* that prepared them for survival on the pathogen, and confered in some unknown way, a degree of specificity to the HSF-1 transcriptional response.

**Fig. 2.**
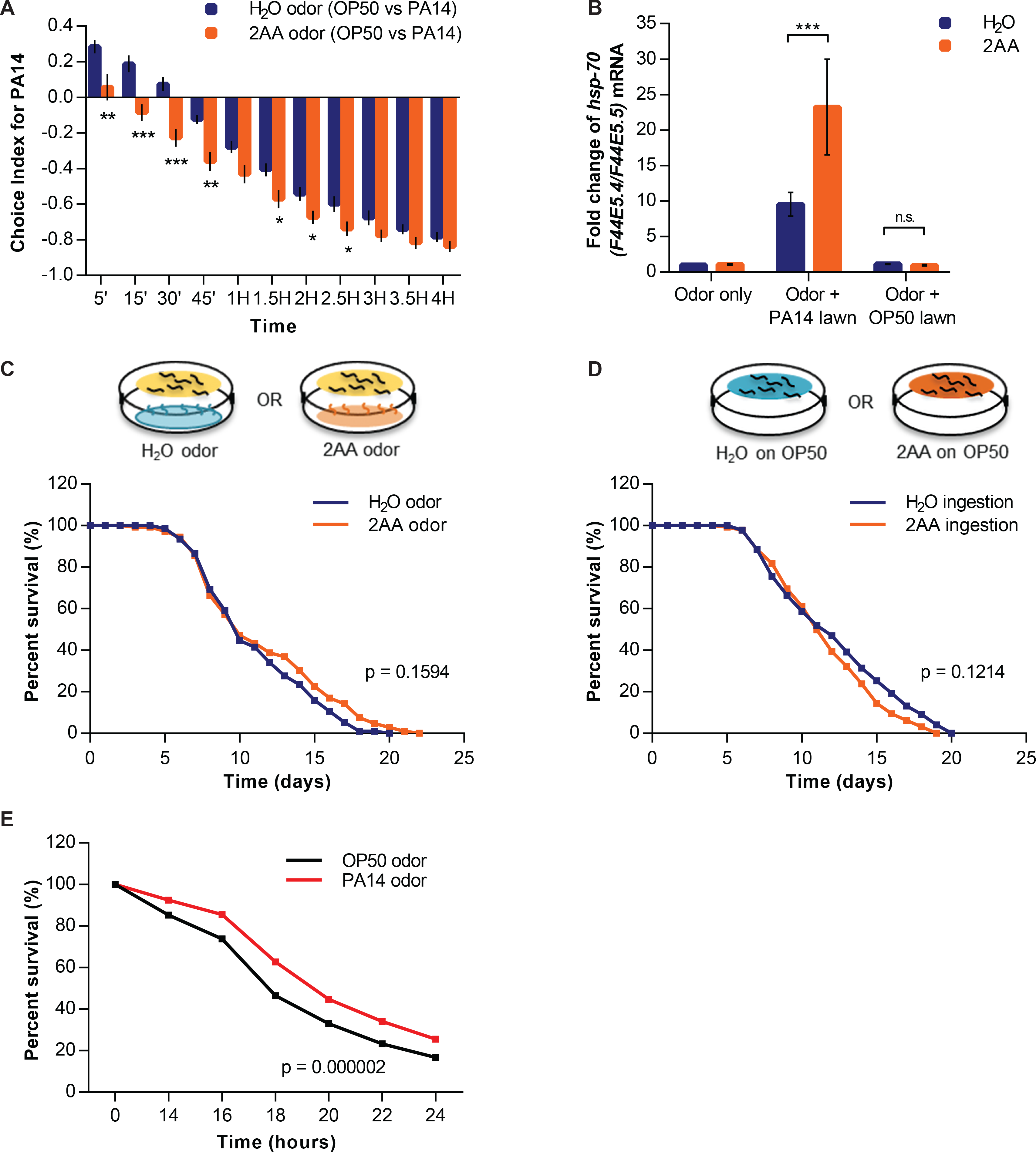
The compound 2-aminoacetophenone (2AA) made by PA14 enhances olfactory avoidance behavior and HSF-1 activation. **(A)** Choice index of wild-type animals for PA14, following pre-exposure to the odor of either water or 2AA. Preference was recorded at the times indicated on the x-axis. N = 10 experiments of 30 animals per condition. Student’s t-test. *p<0.05, **p<0.01, ***p<0.001. **(B)** *hsp-70* (F44E5.4**/**F44E5.5) mRNA abundance measured by qRT-PCR in wild-type animals that were pre-exposed to water ‘odor’ or 2AA odor and subsequently placed on a PA14 or OP50 lawn. Values are relative to animals pre-exposed to water. N = 21 experiments of 30 animals per condition. Pairwise mean comparison from linear mixed model analysis. ***p<0.001. See Materials and Methods and Table S2 for complete details. **(C)** and **(D)** Lifespan curves of animals **(C)** continuously exposed to water (control) or 2AA odor, or **(D)** in physical contact with water-treated (control) or 2AA-treated OP50. N= 3 experiments of 50 animals per condition. Log-rank tests. No significance. See also Table S3 and Materials and Methods. **(E)** Survival of wild-type animals on PA14, following pre-exposure to OP50 or PA14 odor. N = 8 experiments of 50 animals per condition. Log-rank test. p<0.001. Also see Table S4; (A, B) means ± S.E.M, and (C-E) total animals across all experiments. Legends: pre-exposure conditions (choice).

### Serotonin is required for the enhancement of HSF-1-dependent *hsp* gene expression upon olfactory training

In *C. elegans* and other organisms, the neuromodulator 5-HT is known to mediate learning (15, 21, 22, 26, 39-42). We therefore tested whether the enhanced behavioral and transcriptional response to PA14 that occurred following the pre-exposure of *C. elegans* to PA14-derived odors required 5-HT. This was the case. Compared to the 5 minutes needed for wild-type animals trained by PA14 odor to avoid PA14 lawns, animals that lacked functional tryptophan hydroxylase or *tph-1* (43), the rate limiting enzyme for 5-HT synthesis, took 1 hour to avoid PA14 after pre-exposure to the odor of PA14 (Figs. 3A, S3A). This delay in avoidance was indeed due to the lack of 5-HT, as incubation with exogenous 5-HT, a process known to load 5-HT into the serotonergic neurons within a few minutes (42) (Fig. S3B), rescued the deficiency in the learned-response of *tph-1* mutant animals: 5-HT-treated *tph-1* animals trained with PA14 odor acted like wild-type animals trained with PA14 odor and avoided PA14 lawns by 5 minutes (Figs. 3B, S3C; compare to Fig. 1A). For reasons we do not understand, but which may be related to the differential role of 5-HT in modulating innate and learned aversive behavior in other organisms (44), control, mock-trained *tph-1* animals lacking 5-HT avoided PA14 lawns earlier than wild-type control animals (Fig. S3A; compare to Fig. 1A). This aberrant behavior was also reversed with exogenous 5-HT treatment and 5-HT-treated *tph-1* animals mock-trained on OP50 odor acted more like wild-type animals exposed to OP50 odor and avoided PA14 lawns later, by 45 minutes (Figs. S3C, S3D; compare to Fig. 3A). However, this rescue was more variable and did not reach significance.

**Fig. 3.**
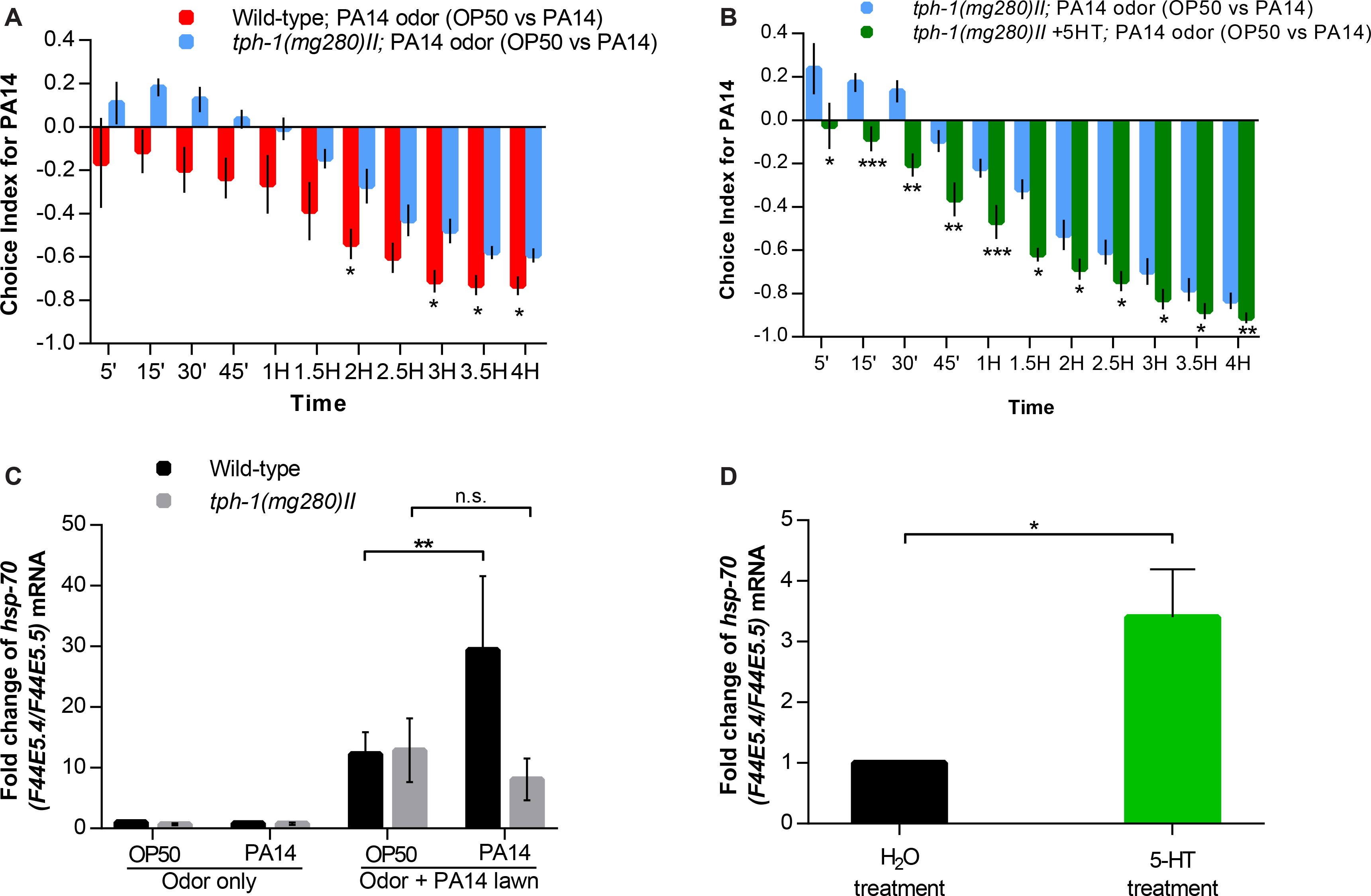
Serotonin is required for olfactory learning-mediated HSF-1 activation. **(A)** Choice index for PA14 of wild-type and *tph-1 (mg280) II* animals pre-exposed to the odor of PA14. Preference was recorded at times indicated (x-axis). N = 3-4 experiments of 30 animals per condition. Student’s two-sample t-test (unequal variance) *p<0.05. **(B)** Choice index for PA14 of *tph-1 (mg280) II* animals pre-exposed to the odor of PA14, compared to the choice indices in *tph-1 (mg280) II* animals treated with exogenous 5-HT (see Fig S3 for rescue), and pre-exposed to PA14 odor. Preference was recorded at times indicated (x-axis). N = 4 experiments of 30 animals per condition. Student’s paired t-test. *p<0.05, **p<0.01, ***p<0.001. **(C)** *hsp-70* (F44E5.4**/**F44E5.5) mRNA abundance measured by qRT-PCR upon PA14 exposure in wild-type and *tph-1 (mg280) II* animals pre-exposed to OP50 or PA14. Values are relative to wild-type animals pre-exposed to the odor of OP50. N = 9 experiments of 30 animals per condition. Pairwise mean comparison from linear mixed model analysis. **p<0.01 for wild-type (odor+PA14 lawn). No significance for *tph-1 (mg280) II* (odor+PA14 lawn). See Materials and Methods and Table S2 for complete details. **(D)** *hsp-70* (F44E5.4**/**F44E5.5) mRNA abundance measured by qRT-PCR following exposure of animals to exogenous 5-HT. Values are relative to control water-treated animals. N = 7 experiments of 20-30 animals per condition. Student’s paired t-test. *p<0.05.

Consistent with the behavioral response, *tph-1* mutant animals were deficient in the enhanced HSF-1-dependent transcriptional response elicited by pre-exposure to PA14 odor (Fig. 3C; Table S2), although 5-HT was not required for HSF-1 activation on PA14 *per se*. This was inferred from the observation that *tph-1* mutant animals did induce *hsp70* (F44E5.4/F44E5.5) expression when exposed to PA14 lawns as assessed by qRT-PCR (Fig. 3C; Table S2). However *tph-1* mutant animals pre-exposed to OP50 or PA14 odors both expressed similar amounts of *hsp70* (F44E5.4/F44E5.5) mRNA upon subsequent encounter with PA14 lawns, and there was no increase in *hsp70* expression based on prior olfactory experience (Fig. 3C; Table S2). We tested whether treatment with exogenous 5-HT could also reverse this defect. However, consistent with what we had previously observed upon optogenetic activation of serotonergic neurons (7), exposure to exogenous 5-HT already induced the expression of *hsp70* (F44E5.4/F44E5.5) mRNA, even without pre-exposure to PA14 odor (Fig. 3D). Although confirming the role of 5-HT in triggering HSF-1 activity, these data suggested that the more nuanced-control over HSF-1-mediated gene expression that occurs within the animal in response to physiological stimuli may be due to a tighter regulation over 5-HT release and availability.

Because 5-HT is synthesized only in neurons in *C. elegans* (39, 43), we tested whether 5-HT-dependent HSF-1 activation was restricted to neurons. This was not the case. Single molecule fluorescent *in situ* hybridization (smFISH) used to detect *hsp70* (F44E5.4/F44E5.5) mRNA across the entire organism indicated that exposure to PA14 induced F44E5.4/F44E5.5 mRNA in all tissue types including neurons, intestine and germline cells, and mRNA expression was enhanced in all these tissues in wild-type animals when trained by PA14 odor (Figs. 4A-F). Consistent with the whole animal qRT-PCR data, *tph-1* mutant animals also induced *hsp70* (F44E5.4/F44E5.5) mRNA when exposed to PA14, but the induction of mRNA in *tph-1* mutant animals remained the same irrespective of prior olfactory training and was also similar to that in control, wild-type animals mock trained with OP50 (Figs. 4A-F). Taken together, these studies showed that both the enhanced avoidance behavior as well as enhanced HSF-1-dependent chaperone gene expression were mediated by the 5-HT learning circuitry.

**Fig. 4.**
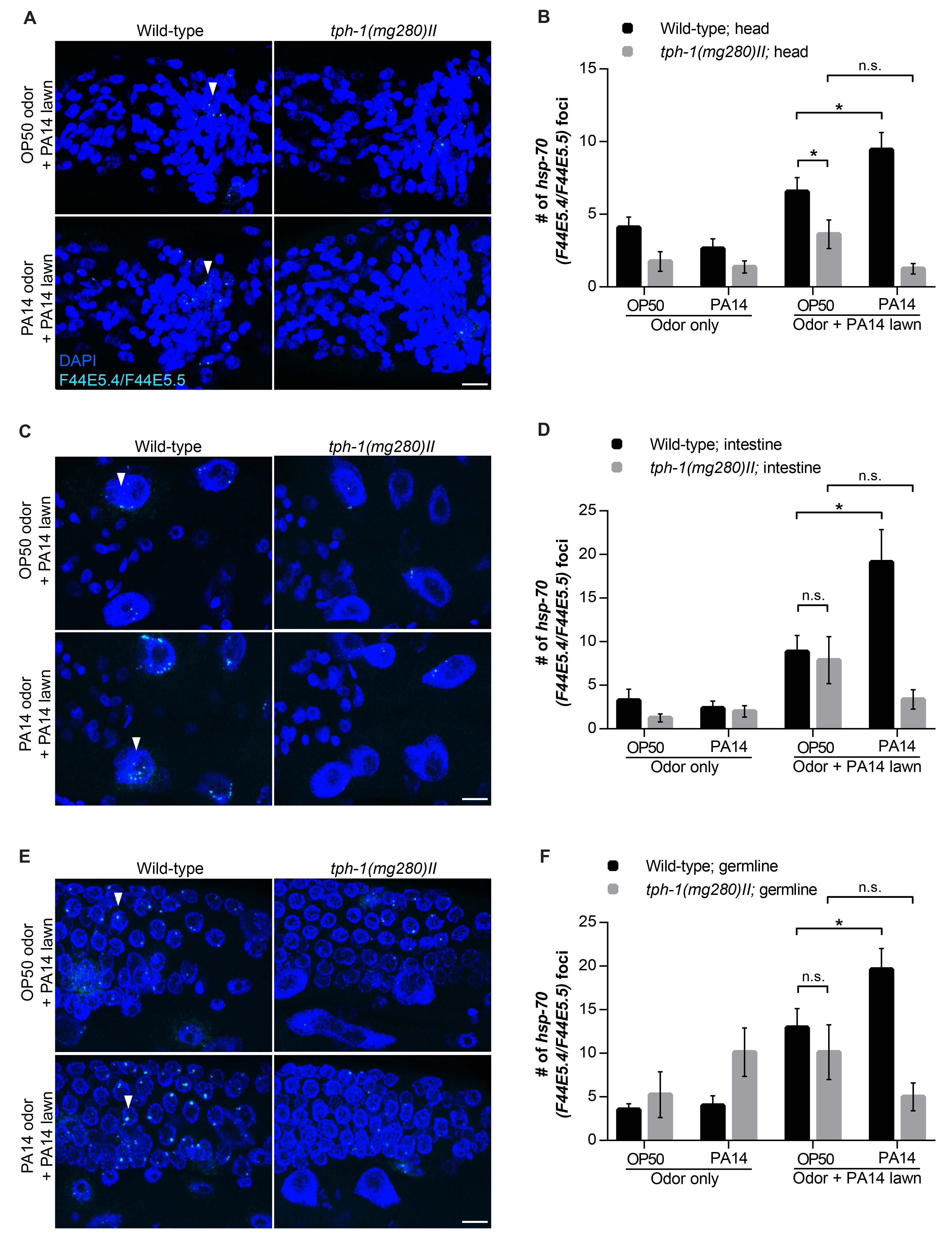
The compound 2-aminoacetophenone (2AA) made by PA14 enhances olfactory avoidance behavior and HSF-1 activation. Serotonin-mediated learning activates HSF-1 throughout the animal. **(A-C)** smFISH confocal micrographs showing *hsp-70* (F44E5.4**/**F44E5.5) mRNA and DAPI in wild-type and *tph-1 (mg280) II* animals pre-exposed to OP50 or PA14, and subsequently subjected to a PA14 lawn. Images are projected Z-stack images of 10 µm sections across the (A) head, (C) intestine and (E) germline of animals. Arrowheads represent *hsp-70* (F44E5.4**/**F44E5.5) mRNA foci. Scale bars, 10μm. **(B, D and F)** Quantification of number of *hsp-70* (F44E5.4**/**F44E5.5) foci, in projected images. N = 8-11 animals per tissue per genotype per condition, quantified from 2-3 independent experiments (see Materials and Methods). Student’s paired t-test, *p<0.05 for wild-type (OP50 odor + PA14 lawn) compared to (PA14 odor + PA14 lawn). No significance for *tph-1 (mg280) II* (OP50 odor + PA14 lawn) compared to (PA14 odor + PA14 lawn). (A, B, D and F) Data represent means ± S.E.M. (A) Legends: genotype; tissue.

### Olfactory training and serotonin localize HSF-1 to nuclear bodies in cells throughout the animal in anticipation of PA14 encounter, priming HSF-1-dependent gene expression

How might olfactory learning enhance HSF-1 transcriptional activity? To answer this, we examined whether olfactory training modified any of the steps known to accompany HSF-1 activation. HSF1-dependent transcription of *hsp* genes is a multistep process that varies to some extent between species (1, 4, 45-50). In mammalian cells, HSF1-dependent *hsp* expression involves conversion of the HSF1 monomers to trimers, increased phosphorylation and other post-translational modifications, acquisition of competence to bind heat shock elements (HSEs) on promoters of *hsp* genes and the recruitment of HSF1 to these HSE elements in a manner that is dependent on the chromatin landscape and transcriptional machinery. We characterized these steps for the transcriptional activation of HSF-1 in *C. elegans* (Fig. S4 and Figs. S5A, B). Consistent with its role as an essential gene in development (51), *C. elegans* HSF-1, as detected by an antibody specific for endogenous *C. elegans* HSF-1 (Figs. S4A, B), and by the localization of a single copy GFP-tagged HSF-1, is constitutively present in nuclei (7, 50, 52) (Fig. S4C) appears to be phosphorylated (Fig. S4D), and is trimerized (Fig. S4E) even at ambient temperatures. Electrophoretic mobility shift assays (EMSA) indicated that in accordance with its trimerization capability at ambient temperatures, *C. elegans* HSF-1 can bind DNA containing canonical *C. elegans* heat shock elements (HSEs) in vitro (Figs. S5A, B). The ability of *C. elegans* HSF-1 to bind HSE-containing DNA in vitro does not change with stress-induced transcriptional activation (Figs. S5A, B). However, in agreement with the lack of expression of inducible *hsp* genes at ambient temperatures in the absence of stress, *C. elegans* HSF-1 did not constitutively bind the *hsp70* promoter region in vivo as assayed by chromatin immunoprecipitation and qPCR (ChIP-qPCR) (Fig. S5C). Instead HSF-1 binding to the *hsp70* (F44E5.4/F44E5.5) promoter in vivo required a stressor such as heat shock which caused transcriptional activation and an approximately five-fold enrichment of HSF-1 at the *hsp70* (F44E5.4/F44E5.5) region (Fig. S5C).

Olfactory pre-exposure to 2AA or PA14 odor did not enhance the ability of HSF-1 to bind DNA in vitro as assayed by EMSA (Figs. S5A, B), nor did it cause HSF-1 to bind *hsp70* promoter regions in vivo as seen by ChIP-qPCR (Fig. S5C). HSF-1 appeared to be phosphorylated upon actual exposure of animals to PA14 lawns, as visualized by its retarded mobility by SDS-PAGE; however, pre-exposure to 2AA did not induce this post-translational change, and HSF-1 in both control (water-exposed) and 2AA-exposed animals appeared identical by SDS-PAGE analysis (Fig. S5D). Exposure to 2AA or PA14 odor alone, however, caused a significant fraction of HSF-1 to re-localize into punctate nuclear bodies (Fig. 5A). This was similar to changes in HSF-1 localization known to occur when HSF-1 is actively transcribing *hsp* genes upon exposure to stressors such as increased temperatures (7, 50, 52) (Fig. S4C), and PA14 lawns (Fig. 5A). Therefore, this formation of nuclear bodies upon 2AA exposure alone was surprising, as exposure to 2AA odor did not induce the transcription of *hsp* genes. The number of nuclei containing HSF-1 nuclear bodies following exposure of animals to the PA14 odorant 2AA, averaged 8.7%, ranging from 3 – 42% amongst the germline nuclei where HSF-1 was the easiest to visualize, and was visible in 71% of animals scored (Figs. 5A, B; N=24 animals and 707 nuclei). In comparison, only an average of 2.2% of germline nuclei ranging between 0-10 % (Figs. 5A, B; N=25 animals, 890 nuclei) of control animals exposed to the odor of water showed any evidence of HSF-1 nuclear bodies. The re-localization of HSF-1 into nuclear bodies was reversible, and the numbers of HSF-1 nuclear bodies in animals exposed to PA14 odorants diminished to control levels (3.0%; N= 17 animals and 633 nuclei) following 30 minutes recovery on OP50, and did not differ from that in control water-exposed animals (3.2%; N= 19 animals and 752 nuclei; Figs. 5A, B).

**Fig. 5.**
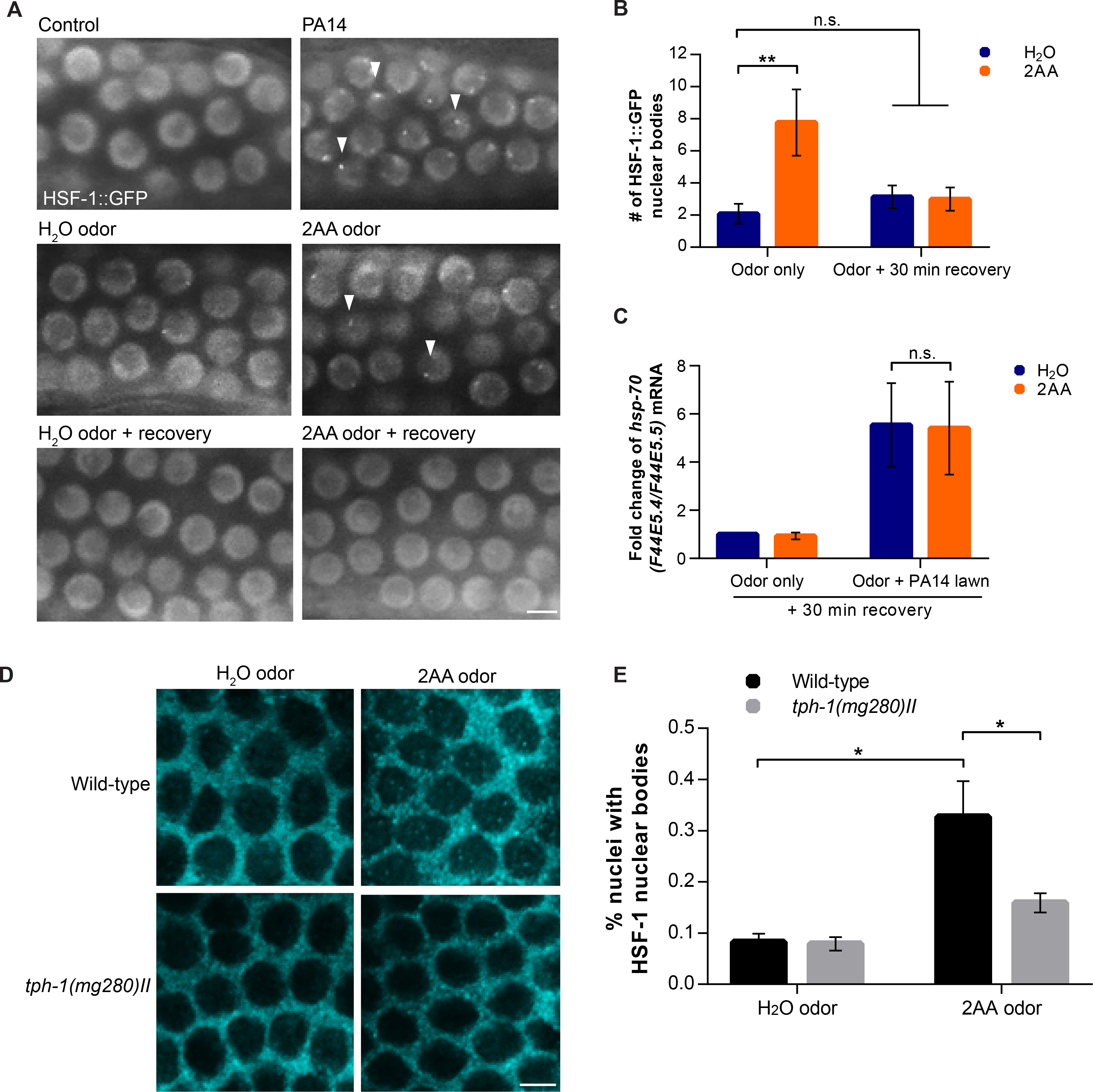
Olfactory learning primes HSF-1 through the formation of HSF-1 nuclear bodies. **(A)** HSF-1::GFP localization in germline nuclei: control animals at ambient temperature, animals on a PA14 lawn, animals pre-exposed to water-odor or animals pre-exposed to 2AA odor, and animals allowed a 30 minute recovery following pre-exposure to water-odor or 2AA odor. Arrowheads indicate HSF-1 nuclear bodies. Scale bar, 5μm. **(B)** Quantification of HSF-1 nuclear bodies under all conditions listed in (A). N= 33-39 nuclei per animal, 17-25 animals per condition, across 3-4 independent experiments (See materials and Methods). Student’s two-sample t-test (unequal variance). **p<0.01 for odor only. No significance for (Odor + 30 min recovery) **(C)** *hsp-70* (F44E5.4**/**F44E5.5) mRNA abundance measured by qRT-PCR following exposure of animals to the odor of water or 2AA and allowed to recover for 30 minutes on OP50 lawn before being placed on PA14 lawns. N=5-9 experiments of 30 animals per experiment. Pairwise mean comparison from linear mixed model analysis. No significance. See Materials and Methods and Table S2 for complete details. **(D)** Confocal micrographs of individual Z-sections showing HSF-1 localization in germline nuclei of wild-type and *tph-1 (mg280) II* animals pre-exposed to water-odor or pre-exposed to 2AA odor. Scale bar, 5μm **(E)** Quantification of HSF-1 nuclear bodies under all conditions listed in (D). N = 50-55 nuclei per animal, 8-18 animals per condition, across 2-3 independent experiments (see Materials and Methods). Student’s two-sample t-test (unequal variance) t-test. *p<0.05.

The HSF-1-mediated transcriptional ‘memory’ of pre-exposure to PA14 odors that resulted in enhanced HSF-1-dependent *hsp* gene expression correlated with the presence of HSF-1 nuclear bodies (Figs. 5A, C). Whereas animals exposed to the PA14 odorant showed the presence of HSF-1 nuclear bodies and displayed enhanced expression of *hsp* genes when placed on PA14 lawns, animals that were allowed to recover for 30 minutes on innocuous OP50 lawns after being trained on PA14 odorants, no longer displayed enhanced *hsp* gene expression when placed on PA14 lawns (Figs. 5A, C; compare with Fig. 1C). In further support of the role of HSF-1 nuclear bodies in the learning-dependent enhancement of HSF-1 transcription, *tph-1* animals that lacked 5-HT, and were deficient in olfactory experience-mediated increase in *hsp* gene expression also had markedly fewer HSF-1 nuclear bodies upon olfactory training (Figs. 5D, E; N=10 animals with 504-545 nuclei per condition). These data, together, indicate that the priming of HSF-1 upon olfactory exposure to PA14 odorants, which resulted in an enhancement of *hsp* gene expression upon actual encounter with PA14 lawns, was occurring through the mobilization of HSF-1 to nuclear bodies throughout cells of the animal.

### HSF-1 nuclear bodies co-localize with RNA polymerase II

Transcription does not occur homogeneously throughout the nucleus, but rather occurs at specialized, discrete sites (53). We hypothesized that because animals that were trained by PA14-derived odor did not induce *hsp70* gene expression unless they encountered PA14 lawns, olfactory signaling may be pre-emptively facilitating the association of HSF-1 with the transcriptional machinery needed to support transcription if the actual encounter with the threat were to occur. We therefore investigated whether the location of the HSF-1 nuclear bodies corresponded to known sites enriched in transcriptional activity. In *C. elegans*, the *hsp-16.2* promoter has been shown to localize to the nuclear pore complex (NPC) following heat shock (53, 54). However, although the HSF-1 nuclear bodies were occasionally in the vicinity of NPCs in germline nuclei, they did not co-localize with NPCs under any conditions (Figs. 6A-C; N = 4-6 animals and 145-180 nuclei per condition). On the other hand, over half of the HSF-1 nuclear bodies (0.2 of the 0.3 nuclear bodies per nucleus, N = 12 animals and 813 nuclei) that were induced by olfactory exposure to 2AA odor co-localized with total RNA polymerase II (Pol II) (Figs. 6D-F). The number of HSF-1 nuclear bodies that co-localized with Pol II remained the same even when HSF-1 was actively involved in Pol II-dependent transcription of *hsp* genes such as upon heat shock, or when animals were exposed to PA14 lawns (Figs. 6D-F). In comparison, only 0.11 of the rare HSF-1 nuclear bodies (0.15 nuclear bodies per nucleus; N = 10 animals and 678 nuclei) visible in control animals co-localized with Pol II (Figs. 6D-F). Consistent with previous reports (45, 55-57), Pol II appeared to cluster in discrete nuclear regions even prior to 2AA exposure or heat shock (Figs. 6D-F). The formation of HSF-1 nuclear bodies, however, did not appear to require Pol II: RNAi-induced knockdown of the large subunit of RNA polymerase II (*ama-1*) substantially decreased the amounts of Pol II protein in oocyte nuclei (Fig. S6A) but did not interfere with the heat-shock induced formation of HSF-1 nuclear bodies in oocytes (Fig. S6B). We conclude from these studies that olfactory training with PA14 odorants was priming HSF-1 by pre-emptively concentrating it at nuclear loci in close proximity to RNA polymerase II. Although we do not yet understand the nature of these nuclear foci where HSF-1 and Pol II were concentrated, or the intracellular mechanisms by which this occurred, taken together, these data suggest that the ability of the serotonin-based learning circuitry to induce the co-localization of HSF-1 with Pol II in nuclei throughout the animal, in anticipation of an impending encounter with PA14 could result in an enhanced transcriptional response upon actual exposure to the pathogen (45, 46, 54-57).

**Fig. 6.**
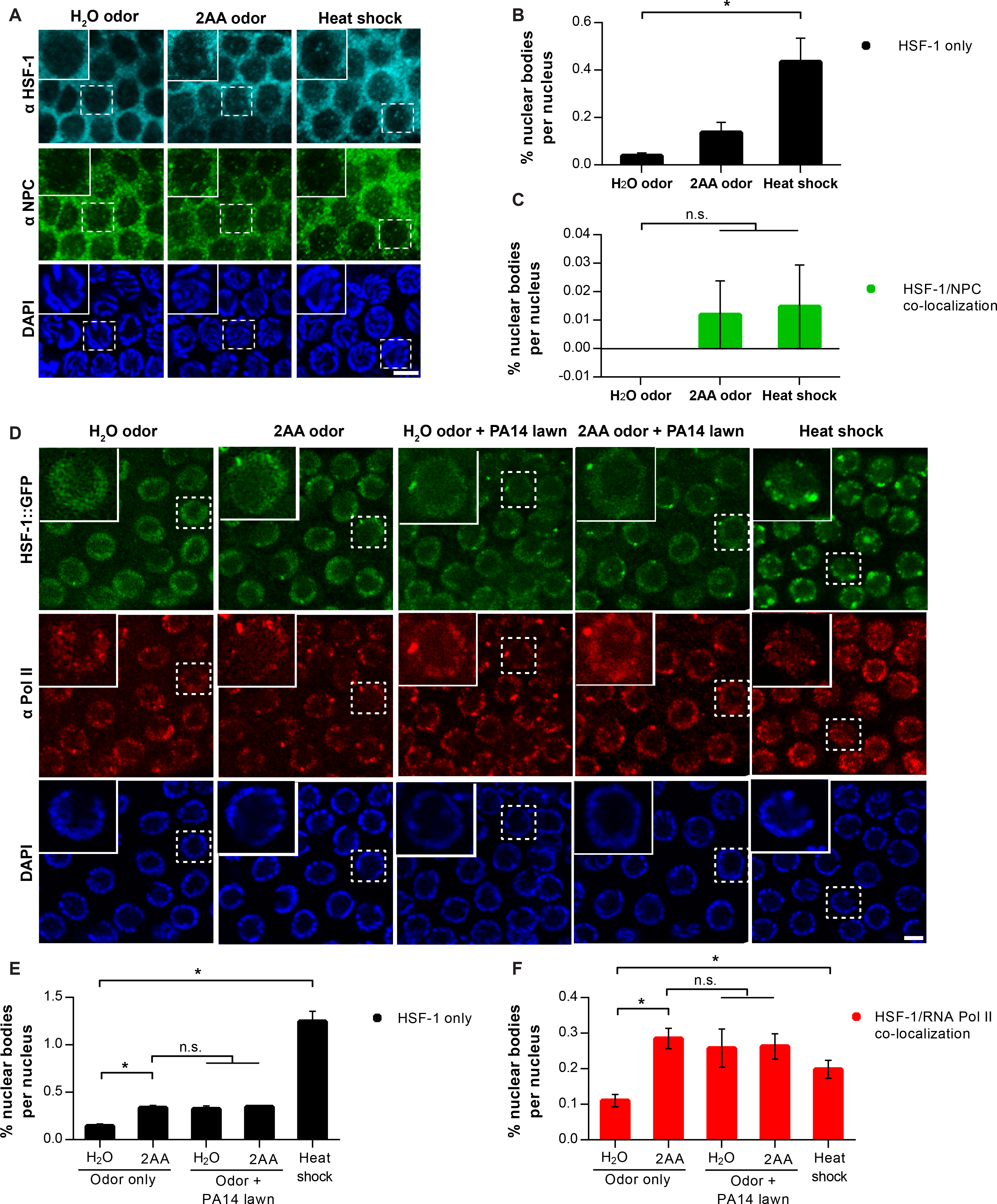
Olfactory learning primes HSF-1 by increasing its association with RNA Polymerase II. **(A)** Immunofluorescence confocal micrographs of HSF-1, Nuclear Pore Complex proteins (NPCs) and DAPI in germline nuclei of wild-type animals exposed to water-odor, 2AA odor and heat shock. Micrographs are individual Z-planes. Scale bar, 5μm. **(B)** Quantification of numbers of total HSF-1 nuclear bodies per nucleus and **(C)** HSF-1 nuclear bodies per nucleus that co-localize with NPCs. N = 30-36 nuclei per animal, 4-6 animals per condition per experiment, 2 independent experiments (see Materials and Methods for details). Student’s two-sample t-test (unequal variance). (B) *p<0.05 when water odor compared to heat shock. (C) No significance between all conditions. **(D)** Immunofluorescence confocal micrographs of HSF-1, RNA Pol II and DAPI in germline nuclei of dissected animals expressing HSF-1::GFP upon exposure to water-odor, 2AA odor, water-odor and PA14 lawns, 2AA odor and PA14 lawns, and heat shock. Micrographs are individual Z-planes. Scale bar, 5μm. **(E)** Quantification of numbers of total HSF-1::GFP nuclear bodies per nucleus and **(F)** HSF-1 nuclear bodies per nucleus that co-localize with RNA Pol II in (D). N =number of nuclear bodies per nucleus in 68-82 nuclei per animal, 8-12 animals per condition per experiment, 3 independent experiments. Student’s two-sample t-test (unequal variance). *p<0.05. (B, D and F) Data represent means ± S.E.M.

### HSF-1 is required for the learned avoidance behavior of C. elegans towards PA14

Not only was HSF-1 activity enhanced by aversive olfactory stimuli, HSF-1 appeared to be required for the behavioral avoidance of PA14. Decreasing the amounts of *hsf-1* mRNA and protein (Fig. S4A), abrogated the behavioral plasticity observed upon exposure to PA14 and animals deficient for *hsf-1* remained equally distributed between the PA14 and OP50 lawns displaying a marked deficiency in their avoidance of PA14 (Figs. 7A, S7A). Loss of *hsf-1* slightly retards motility, causing a delay of ~102 seconds for *hsf-1* RNAi treated animals to traverse the 1 inch distance between the PA14 and OP50 lawns when compared to wild-type animals (Fig. S7B). However, this slight decrease in motility rates could not account for the lack of avoidance behavior of *hsf-1* RNAi treated animals, as they did not avoid PA14 even by 4 hours after being given the choice between PA14 and OP50. By this time, all wild-type animals raised on control RNAi, whether trained on PA14 or control-RNAi odors, had left the PA14 lawns (Fig. 7A). It therefore appeared that 5-HT signaling was integrating olfactory information and HSF-1 activation to flag a sensory stimulus as a threat, providing a basis for the coupling of the enhanced behavioral aversion with the enhanced transcriptional response seen in our experiments. To test if this was the case we activated 5-HT release using optogenetic methods while co-incidentally exposing animals to the odor of the attractive *E. coli* HT115 (Fig. S7C). We predicted that although HT115 does not activate HSF-1 or evoke an avoidance response on its own, optogenetically activating serotonergic neurons so as to activate HSF-1 (7, 58) in the presence of HT115 odor, may change the valence of HT115 from that of attraction to one of aversion, and animals would now avoid HT115. This was the case (Figs. 7B, C). The optogenetic activation of 5-HT release in the presence of HT115 odorant elicited a transient aversive response of animals from HT115, and this aversion lasted for as long as 45 minutes following stimulation when animals were given a choice between HT115 and PA14. Control animals that were mock stimulated by light did not change their behavior and, as expected, were attracted to HT115 and repelled by PA14 (Figs. 7B, C). Thus, inducing 5-HT release during an olfactory stimulus appeared to be sufficient to associate olfactory information regarding odor with HSF-1 activation to trigger an aversive response of *C. elegans* to danger.

**Fig. 7.**
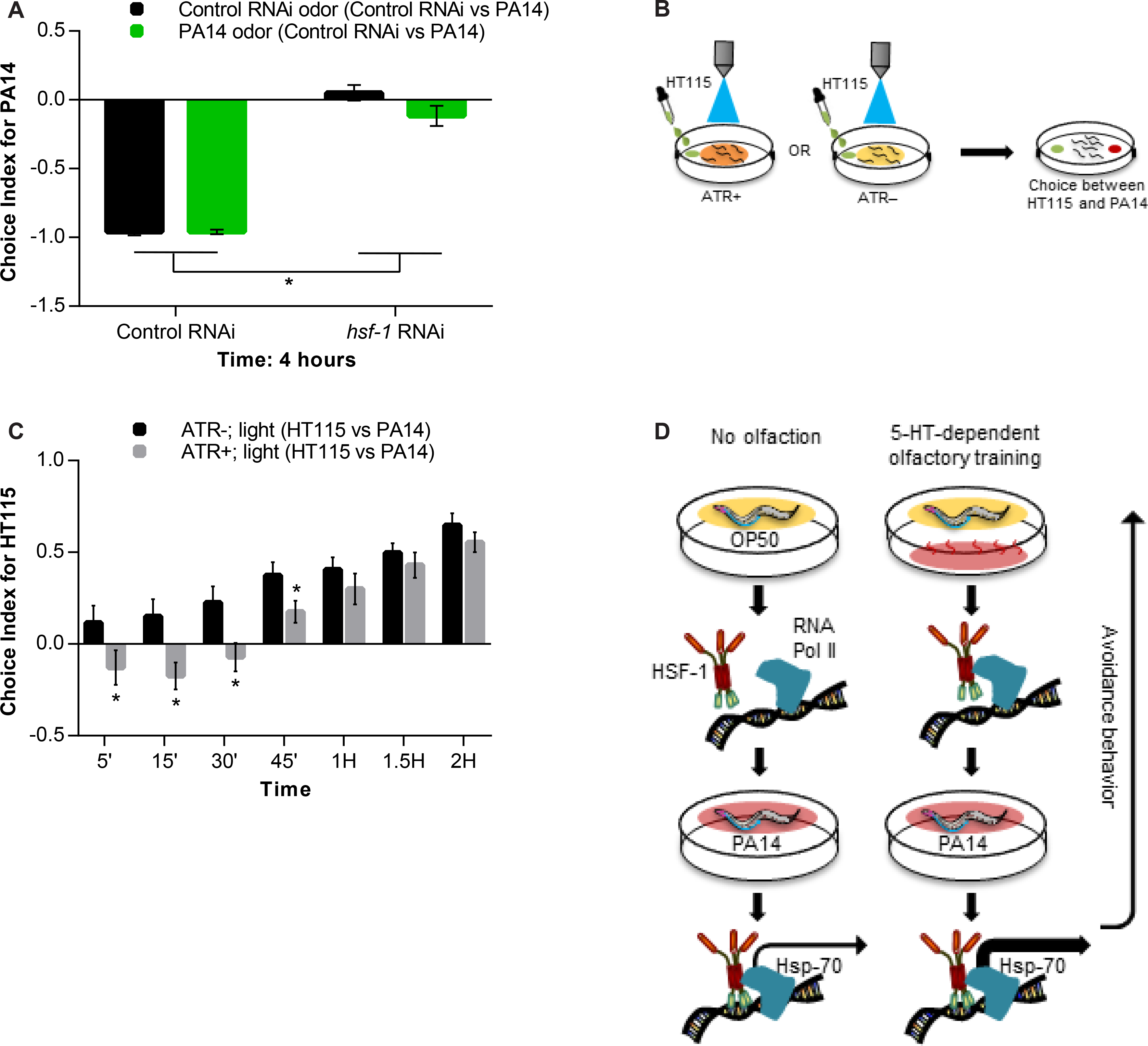
5-HT signaling couples olfactory information with HSF-1 activation to mark sensory stimuli as threats. **(A)** Choice index for PA14 of wild-type animals subjected to control RNAi (empty vector), or *hsf-1* RNAi. Preference was recorded at time indicated (x-axis). N = 5-6 experiments of 30 animals per condition. Student’s two-sample t-test (unequal variance). *p<0.05. **(B)** Schematic of olfactory pre-exposure to HT115 odor in conjunction with optogenetic excitation of serotonergic neurons, followed by behavioral choice assay. ATR+ indicates the presence of all-*trans* retinal required for the light-induced excitation of channelrhodopsin (expressed in the serotonergic neurons) necessary to release 5-HT. ATR– indicates control-mock excited animals. **(C)** Choice index for HT115 in animals +/- ATR following optogenetic excitation of serotonergic neurons. N = 6 experiments in triplicate of 10 animals per condition. Student’s paired t-test *p<0.05. **(D)** Model: 5-HT-dependent olfactory learning facilitates the association between RNA Pol II and HSF-1, resulting in enhanced avoidance behavior as well as enhanced transcription of HSF-1 targets in a stressor-specific manner. (A and C) Data represent means ± S.E.M. (A and C) Legends: pre-exposure condition (choice).

## Discussion

In summary, our data provide a mechanism whereby 5-HT-dependent learning and HSF-1 activation are coupled to elicit behavioral avoidance and transcription of cytoprotective chaperone genes under threat, enhancing the survival of the animal (Fig. 7D). Our data suggest that 5-HT release from neurons needs to be reinforced by HSF-1 activation throughout the animal to interpret a signal as aversive. Conversely, HSF-1 itself is activated by 5-HT release in a multi-step process. In our experiments we show this in some detail, in response to the odorants of the toxic bacteria *Pseudomonas*. However, similar responses could underlie the reaction of *C. elegans* to other stressors. The neuroethological significance of the response of *C. elegans* to 2AA seen in our studies is unknown. Our data suggest that 2AA acts as a kairomone (59)- an interspecies chemical messenger whose adaptive benefit appears to be greater for the recipient than the emitter. *C. elegans* is a bacterivore and relies, like its related parasitic nematode species, on chemical cues to interpret the hostility or hospitality of its environment. However, because 2AA in itself does not elicit an aversive response suggesting that it is less like a predator odor or danger pheromone, we speculate that it is akin to what a loud noise may signify to a human- a reason for investigation, to be coupled with an avoidance response if confirmed to be associated with danger. 2AA is also secreted by other known pathogens of *C. elegans* such as *Burkholderia sp*. and arthropods (60-62), perhaps accounting for the ability of *C. elegans* to detect it and to effectively modulate its behavior and stress responsiveness.

Our data also suggest that neuronal control over HSF-1-dependent transcription of chaperones genes within the metazoan *C. elegans* is at least a two-step process. The first step, the reversible and anticipatory change in nuclear localization of HSF-1, mediated by neurons and 5-HT, pre-emptively promotes HSF-1 concentration at nuclear regions close to RNA polymerase II, and could conceivably prepare the chromatin and transcriptional machinery for transcription, were the stressor to materialize. This could allow animals to enhance chaperone gene expression upon encounter with the actual stressor (45, 46, 54-57) (Fig. 7D). Nevertheless, a subsequent, as yet unknown signal ‘confirming’ the threat, also likely dependent on 5-HT, appears to be required for the actual transcription of *hsp* genes (Fig. 7D). This signal confers the specificity of the transcriptional response to the stressor. The exact mechanism by which 5-HT dependent learning induces HSF-1 to organize into nuclear bodies, and the nature of these structures and the genomic regions in the vicinity (63, 64) remain to be investigated. In *Drosophila* and mammalian cells, a fraction of RNA polymerase II is held paused at *hsp* loci until HSF-1 binding initiates transcription and the release of Pol II into the gene body (45, 47, 56, 65, 66). However, consistent with our data, in these cells too HSF-1 binding alone is not the determining event for the release of Pol II pausing, as HSF-1 can bind *hsp70* loci without inducing transcription (67).

The multistep activation of a fundamental cytoprotective response to a threat raises intriguing questions. Given its extraordinarily beneficial roles in conferring stress-resistance, why not simply activate HSF-1 in anticipation, even upon the slightest hint of stress? We believe that the answer to this may lie in findings that high levels of chaperone expression disrupt basic functions of a cell such as growth, division and secretory functions, and increase susceptibility to transformation (68-70). In fact, it has been shown that chaperone levels within cells of a multicellular organism are not maintained in excess (71). We hypothesize, therefore that for cells integrated into a metazoan, activation of HSF-1 needs to be tightly controlled to occur only upon confirmation of danger, so as to prevent the possible disruption of tissue homeostasis. Organisms survive a range of environmental fluctuations and have evolved to colonize a vast diversity of environmental niches despite the sensitivity of protein-based biological processes to environmental perturbations. We believe that our data begin to address one mechanism through which such adaptation could occur.

## Materials and Methods

### C. elegans Strains

The following *C. elegans* strains were used. Generation of the AM1061 transgenic strain is described in Tatum *et al*, 2015(7). The remaining strains were obtained from the Caenorhabditis Genetics Center (CGC).

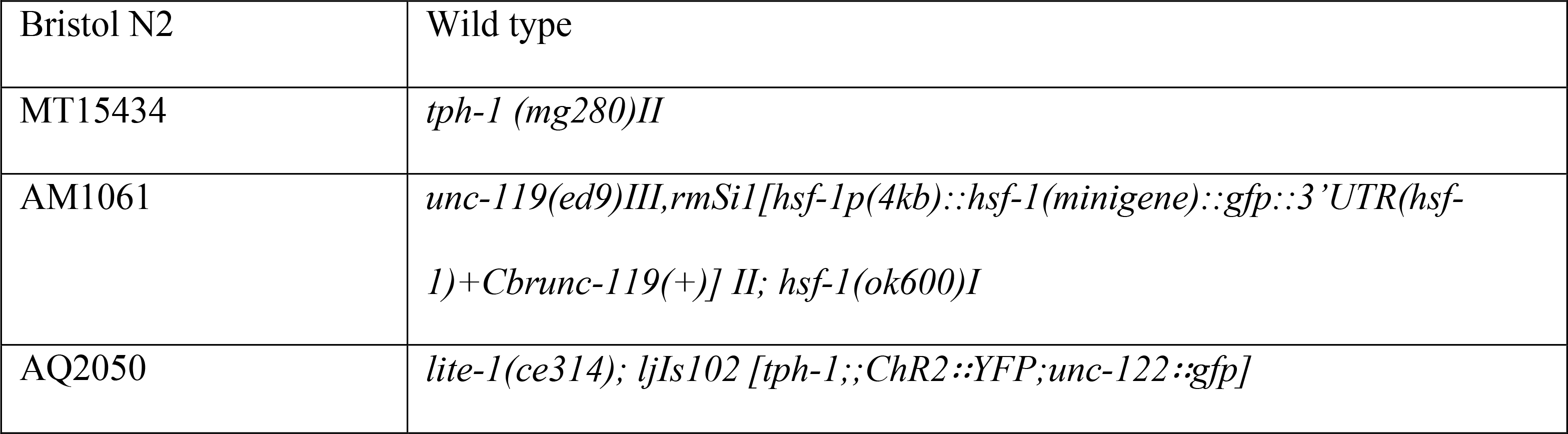

### Growth conditions of *C. elegans* and bacteria

All strains were grown and maintained at 20°C. Ambient temperature was maintained at 20-22°C, and carefully monitored throughout the experimental procedures. All animals included in the experiments, unless stated otherwise, were one day old hermaphrodites that were staged as L4 animals 24-26 hours prior to the start of the experiment. Worms were grown and maintained at low densities, under standard conditions in standard incubators (20°C), as previously described (7). Specifically, animals were fed with OP50 obtained from the Caenorhabditis Genetics Center (CGC, Twin Cities, MN) that were seeded 2 days prior to use, and stock strains were maintained at low densities by passaging 8 or 10 L4s onto NGM plates, and 4 days later picking L4 animals onto fresh plates for experiments. The NGM plates were standardized by pouring 8.9ml liquid NGM per plate that yielded plates with an average weight of 13.5 ± 0.2 g. Any plates that varied from these measurements were discarded. The *Pseudomonas aeruginosa* strain PA14 was obtained from the Yahr Lab (University of Iowa) and the *Serratia marcescens* strain DB11 was obtained from the CGC. Both PA14 and DB11 lawns were kept at 25°C for two days prior to use in experiments.

### RNA Interference (RNAi) methods and Verification of RNAi-induced knock-down

RNAi experiments were conducted using the standard feeding RNAi method (72-74). Bacterial clones expressing the control (empty vector) construct and the ds RNA targeting the majority of *C. elegans* genes were obtained from the Arhinger RNAi library (72) now available through Source Bioscience (http://www.us.lifesciences.sourcebioscience.com/clone-products/non-mammalian/c-elegans/c-elegans366-orf-rnai-library-v11/). The RNAi clones used in experiments were sequenced for verification prior to use. The pL4440 empty vector was used as control RNAi. RNAi induced knockdown was conducted by either feeding animals for 24 hours (*ama-1*) or for over one generation, where 2^nd^ generation animals were born and raised on RNAi bacterial lawns (*hsf-1*). RNAi-mediated knockdown was confirmed by scoring for known, knock-phenotypes of the animals subject to RNAi that have been reported in genome-wide RNAi screens in *C. elegans* (slow and arrested larval growth as well as larval arrest at 27ºC for *hsf-1* RNAi, and 2^nd^ generation embryonic lethality in the case of the *ama-1* RNAi). Knock-down was further ascertained using either western blots (HSF-1) or immunofluorescence (AMA-1) to verify a decrease in protein levels.

### Olfactory pre-exposure

Bacterial cultures were grown in LB broth to OD600 values of between 1.4-1.7, and the variation between cultures within an experiment was kept to ± 0.1. For pre-exposure to bacterial odors, experiments were carried out in a 25°C incubator, and 750ul of culture was placed in the lid of a 35x10mm petri dish (VWR International, Catalog # 10799-192; Radnor, PA), which was then placed in the lid of an inverted standard NGM petri dish (see image below). As the plates were inverted, animals crawled on OP50 lawns on the ‘top’ of the plates, while the odorant remained on the ‘bottom’, undisturbed, and so, at no point, did animals come in contact with the odorant. L4 animals were picked onto these NGM plates on OP50 lawns on the previous day, and remained on their respective OP50 lawns during the course of the exposure to odor. We verified that no bacterial spores were transferred via this exposure by conducting the same procedure with an unseeded NGM plate, and observing the plates over the course of the next two days for bacterial growth. For “naïve” conditions, the animals were not given any odor prior to the start of the experiments. When the pre-exposure was to water or 2AA (Catalog no. A38207; Sigma-Aldrich, St. Louis, MO), 3 mls of water or 1mM 2AA (kept at 37°C for 5 minutes prior to use) was used in place of the bacterial culture, and experiments were carried out at room temperature, ranging from 20-22°C. When the pre-exposure was to ethanol or N-(3-Oxododecanoyl)-L-homoserine (3OC12; Catalog no. O9139; Sigma-Aldrich, St. Louis, MO), 3 mls of 0.2% ethanol or 10µM 3OC12 was used and experiments were carried out at room temperature. For experiments with a recovery condition, after the 30 minutes of olfaction, the plate containing the liquid odorant was removed and animals were allowed to “recover” at room temperature for 30 minutes prior to harvesting for subsequent experiments.

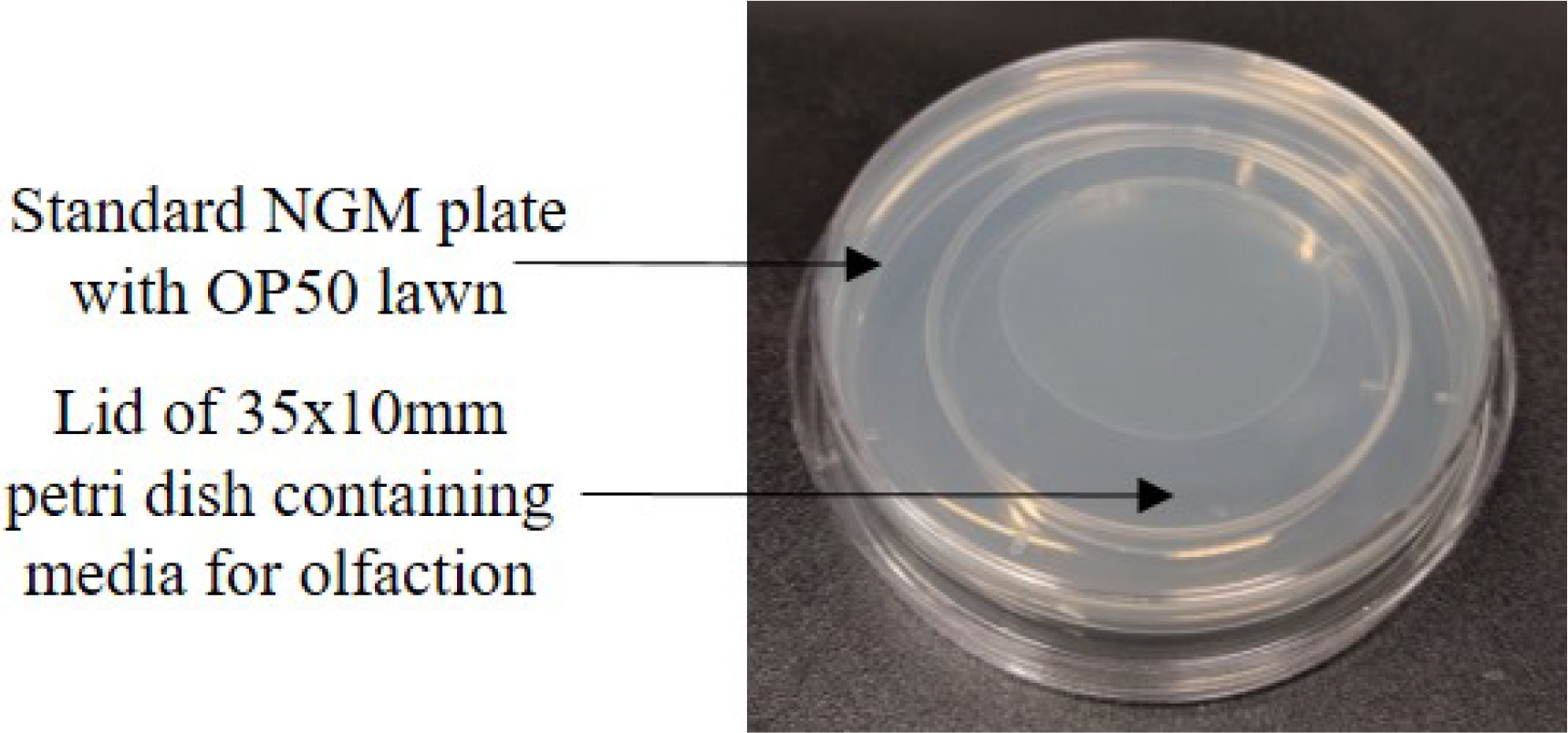

### Bacterial lawn choice assays

L4s were harvested 24-26 hours prior to the start of the behavioral choice experiments. Choice plates were seeded and grown at 25°C for two days before use. For PA14 lawns the duration of growth on the NGM plates prior to the behavioral assay was particularly important. Younger lawns elicited later avoidance behaviors. 85ul of bacterial culture (OP50, PA14, HT115 or DB11) grown to OD600 values of between 1.3-1.6 was used for seeding each lawn. The distance between the lawns was 1 inch (see cartoon below). Following olfactory training (conducted as described above), 30 day one adult animals were transferred to the middle of the choice plates, at a point equidistant from the middle of each lawn and the behavior of the animals was observed at said frequencies for the next 4 hours at room temperature. The number of animals present on either bacterial lawns or off bacterial lawns was recorded, and the experimental bacteria choice index was calculated using the following equation:

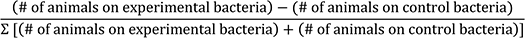

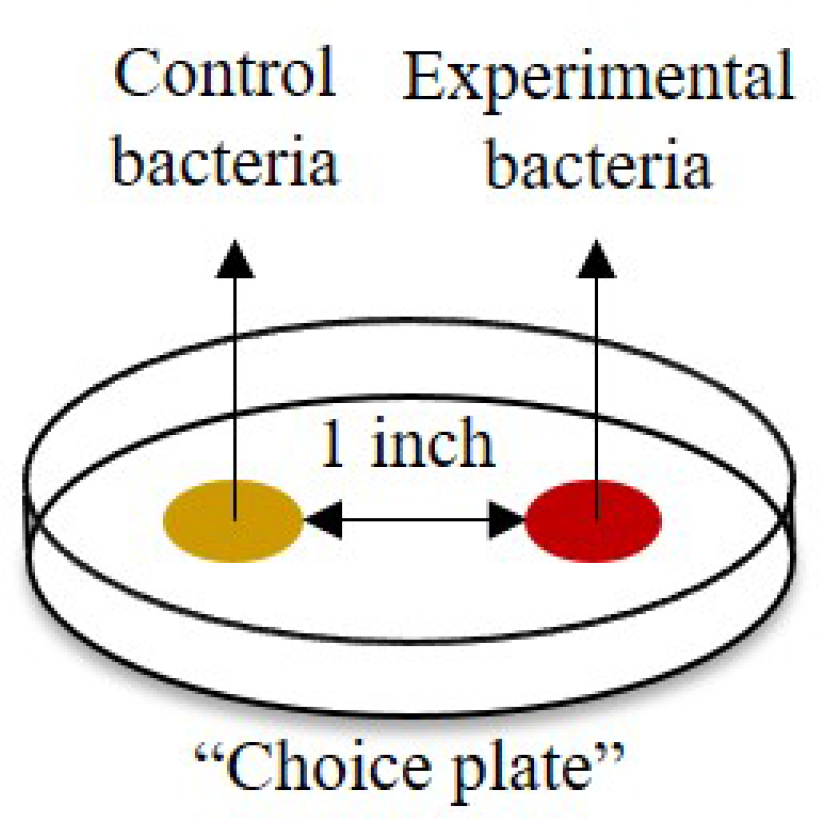

Animals were considered to be on a lawn as long as they were physically in the lawn be it on the edge or in the middle of the lawn at the time of observation at a maximum of 4.0x magnification.

### Chemotaxis assays

Chemotaxis between water and 10mM 2AA were carried out at room temperature. 5ul each of water and 2AA (pre-mixed with 0.5M sodium azide - Catalog no. S2002; Sigma-Aldrich, St. Louis, MO – such that the final concentration of 2AA was 10mM and that of sodium azide was 0.25M) were dropped onto an unseeded NGM plate. The two spots were 1.5 inches apart from each other. The spots were air-dried for 5 minutes, and then the chemotaxis assay was carried out by placing 30 day one worms (harvested as L4s the day before) at a point on the plate equidistant from the two spots. Worms were counted as having made their choice only if they were immobilized at the time points at which observations were made. Choice Index for PA14 was calculated as:

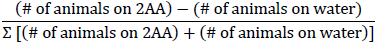

### Exogenous serotonin (5-HT) treatment

Exogenous serotonin treatment was modified from Jafari *et al*, 2011. A serotonin solution (Catalog no. 85036; Sigma-Aldrich, St. Louis, MO) in sterile water was dropped onto the surface of OP50 bacterial lawns (such that the lawns were fully covered in serotonin) on NGM plates and dried for ~2 hours at room temperature prior to use. For confirmation of serotonin uptake using immunofluorescence, serotonin concentrations between 2-20mM were used and day one animals were placed onto serotonin-treated plates for between 30 minutes to 2 hours. For the exogenous serotonin choice assays, a 2mM serotonin solution was used and L4s were then picked onto these plates and kept at 20°C for 24-26 hours prior to experimenting with these day one adults the next day. Olfactory pre-exposure was carried out using OP50 and PA14 bacterial cultures as described above, followed by choice assays as described above.

### PA14 Survival Assays

Survival assays were carried out on day one adult worms that had been harvested as L4s the previous day (50 worms per plate). PA14 killing was performed in liquid bacterial culture in 6-well dishes(75). OP50 liquid bacterial culture used as a control. Both PA14 and OP50 bacteria were grown to an OD600 range of 0.8 – 1.0. Olfactory pre-exposure was performed as described above using OP50 and PA14, and immediately after olfaction, worms were picked into liquid bacterial culture in the 6-well dishes. Plates were covered loosely to allow for air circulation and kept in a 25°C incubator set to 85rpm, and scored periodically for survival by visualization under a microscope. Animals were scored as dead if they were not moving in response to gentle swirling of the media and if there was no pharyngeal pumping. At the end of the experiment, death was confirmed by pipetting the animals onto unseeded plates and lack of revival.

### Longevity Assays

Each experiment was carried out on 50 day one adults harvested as L4s the prior day. For longevity with olfaction, animals were pre-exposed to water and 2AA as described above, and then transferred onto a new OP50-seeded NGM plate. Animals were transferred every two days to avoid starvation, until the point where they were no longer capable of reproduction, typically at day 9. The 2AA liquid, and water, were refreshed on a daily basis. For longevity with ingestion, water and 1mM 2AA was dropped onto OP50 lawns on NGM plates and allowed to dry for 2 hours prior to use. Animals were transferred every two days until day 9, and the water and 2AA-treated plates were made fresh on the day of use. Animals were scored as dead if they were not moving in response to tapping of the plate, or a gentle touch on the NGM adjacent to the animal. Animals that died of internal hatching were discarded.

### Thermotolerance Assays

These assays were carried out on day one adult worms that had been harvested as L4s the previous day, with 20 worms per plate. Olfactory pre-exposure was performed as described above, and immediately after olfaction, worms were subjected to an extended heat treatment (45 minutes) in a circulating water bath pre-heated to 37.5°C. After this heat exposure the animals were allowed to recover for 16 hours at 20°C and were then scored as live or dead the following day. The lack of pharyngeal pumping and lack response to gentle and harsh touch were the criteria used for scoring an animal as dead.

### RNA extraction and quantitative RT-PCR

Samples for RNA were day one adult worms that had been harvested as L4s the previous day, with 30 worms per plate. Olfactory pre-exposure was performed as described above, and animals were either immediately harvested (olfaction only) or subjected to a PA14 lawn for 10 minutes (olfaction + lawn) and then harvested. RNA extraction was conducted according to previously published methods (6, 7). RNA samples were harvested in 50μl of Trizol (Catalog no. 400753; Life Technologies, Carlsbad, CA) and snap-frozen immediately in liquid nitrogen. The following steps were either carried out immediately after snap-freezing or samples were stored at −80°C. Samples were thawed on ice and 200μl of Trizol was added, followed by brief vortexing at room temperature. Samples were then vortexed at 4°C for at least 45 minutes to lyse worms completely. RNA was then purified as detailed in the manufacturer’s protocol with appropriate volumes of reagents modified to 250μl Trizol. RNA pellet was dissolved in 17μl RNase-free water. RNA was treated with DNase using TURBO DNA-free kit (Catalog no. AM1907; Life Technologies, Carlsbad, CA) as per manufacturer’s protocol. cDNA was generated by using the iScript™ cDNA Synthesis Kit (Catalog no. 170-8891; Bio-Rad, Hercules, CA). Real-time PCR was performed using LightCycler^®^ 480 SYBR Green I Master Mix (Catalog no. 04887352001; Roche, Basel, Switzerland), in LightCycler^®^ 480 (Roche, Basel, Switzerland) at a 10μl sample volume, in a 96 well white plate (Catalog no. 04729692001; Roche, Basel, Switzerland). The relative amounts of *hsp* mRNA were determined using the “Delta Delta CT” (ddCT) Method for quantitation. *act-1*, *syp-1* and/or *pmp-3* mRNA was used as internal controls. The use of *syp-1*, which is a germline expressed gene controlled for the variable numbers of embryos that were in the animals when they were prepared for mRNA extraction. All relative changes of *hsp* mRNA were normalized to either that of the wild type control, or the control for each genotype (specified in figure legends). CT values were obtained in triplicate for each sample (technical triplicates). Each experiment was then repeated a minimum of three times. For qPCR reactions, the amplification of a single product with no primer dimers was confirmed by melt-curve analysis performed at the end of the reaction. No reverse transcriptase (no-RT) controls were included to exclude any possible genomic DNA amplification. Primers were designed using Roche’s Universal ProbeLibrary Assay Design Center software and generated by Integrated DNA Technologies (Coralville, IA). The primers used for the PCR analysis were:

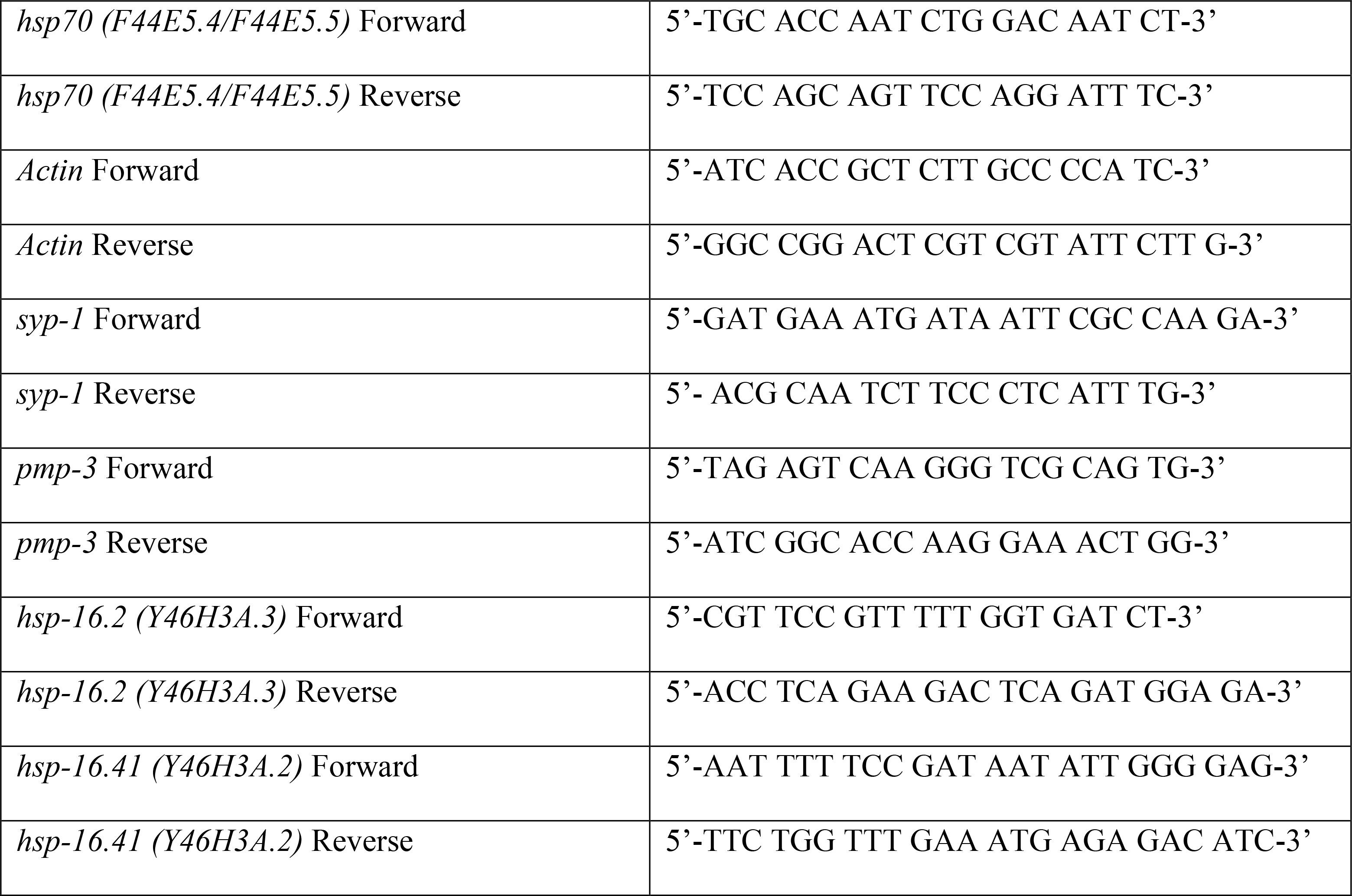

### Single molecule fluorescent in situ hybridization (smFISH)

smFISH probes were designed against F44E5.4/5 by utilizing the Stellaris FISH Probe Designer (Biosearch Technologies, Inc., Petaluma, CA) available online at www.biosearchtech/com/stellarisdesigner. The fixed worms were hybridized with the F44E5.4/5 Stellaris FISH Probe set labeled with Cy5 dye (Biosearch Technologies, Inc., Petaluma, CA), following the manufacturer’s instructions available online at www.biosearchtech.com/stellarisprotocols. 10-20 day one adult wild-type or *tph-1(mg280)II* worms per condition (30’ OP50 olfaction, 30’ PA14 olfaction, 30’ OP50 olfaction + 10’ PA14 lawn, 30’ PA14 olfaction + 10’ PA14 lawn) were harvested, washed once in 1X RNase-free PBS (Ambion, Catalog # AM9624), fixed in 4% paraformaldehyde and subsequently washed in 70% ethanol at 4°C for approximately 24 hours to permeabilize the animals. Samples were washed using Stellaris Wash Buffer A (Catalog # SMF-WA1-60, Biosearch Technologies, Inc., Petaluma, CA) and then the hybridization solution (Catalog # SMF-HB1-10, Biosearch Technologies, Inc., Petaluma, CA) containing the probes was added. The samples were hybridized at 37 °C for 16 hours, after which they were washed three times with Wash Buffer A, then incubated for 30 minutes in Wash Buffer A with DAPI. Following DAPI staining, worms were washed with Wash Buffer B (Catalog # SMF-WB1-20, Biosearch Technologies, Inc., Petaluma, CA) and mounted on slides in approximately 12 μl of Vectashield mounting medium (Catalog # H-1000; Vector Laboratories, Burlingame, CA). Imaging of slides was performed using a Leica TCS SPE Confocal Microscope (Leica, Solms, Germany) using a 63x oil objective. LAS AF software (Leica, Solms, Germany) was used to obtain and view z-stacks and quantification was conducted visually by counting the number of F44E5.4/5 puncta present in nuclei in the head, intestine and germline of each individual worm.

### Western blot protein analysis

For all western blot analyses animals were day one adults. Acute heat shock was performed by wrapping NGM plates with parafilm and sealing plates in a zippered plastic bag. Plates were submerged in a circulating water bath set to 34°C for 10 minutes. For protein analysis, 30 day one adult worms were collected into 18 μl of 1X PBS (pH 7.4) and then 4X Laemmli sample buffer (Catalog #1610737, Bio-Rad, Hercules, CA) supplemented with 10% beta-mercaptoethanol was added to each sample before boiling for 30 minutes. Whole worm lysates were resolved on 8% SDS-PAGE gels and transferred onto nitrocellulose membrane (Catalog # 1620115, Bio-Rad, Hercules, CA). Immunoblots were imaged using Li-Cor^®^ Odyssey Infrared Imaging System (LI-COR Biotechnology, Lincoln, NE). Rabbit anti-HSF1 primary antibody (Sigma Catalog # HPA008888) was used to detect HSF-1 while the mouse anti-alpha tubulin primary antibody (AA4.3), developed by Walsh, C., was obtained from the Developmental Studies Hybridoma Bank, created by the NICHD of the NIH and maintained at The University of Iowa, Department of Biology, Iowa City, IA 52242. The following secondary antibodies were used: Sheep anti-Mouse IgG (H&L) Antibody IRDye800CW^®^ Conjugated (Catalog # 610-631-002, Rockland antibodies & assays, Limerick, PA), Alexa Fluor 680 goat anti-rabbit IgG (H+L) (Catalog # A21109, Molecular Probes, Invitrogen, Carlsbad, CA). The Li-Cor Image Studio software was used to quantify protein levels in different samples, relative to alpha-tubulin levels. Subsequent analysis of protein levels was calculated relative to wild-type controls. For Phos-tage PAGE analysis, Phos-tag reagent was obtained from Wako Pure Chemicals Industries Ltd (Japan) and the protocol was provided here http://www.wako-chem.co.jp/english/labchem/journals/phos-tag_GB2013/pdf/Phos-tag.pdf. Our experiments utilized 25uM final concentration of phos-tag reagent in a 6% SDS-PAGE gel. Full-length recombinant *C. elegans* HSF-1 protein was a kind gift of Dr. Richard Morimoto (Northwestern University, Chicago, IL). For EGS cross-linking experiments, whole worm lysate was prepared by washing worms off the plate in lysis buffer (10mM HEPES pH 7.4, 130mM NaCl, 5mM KCl, 1mM EDTA, 10% glycerol) supplemented with 1mM DTT, 0.2% NP-40 and protease inhibitor cocktail (ThermoFisher Catalog # 87785). Worms were lysed in a Precellys24 homogenizer (Bertin Corp., Rockville, MD) with VK05 beads (Bertin Corp., Rockville, MD) and cleared lysate was incubated at room temperature with 0, 0.1 or 0.5mM EGS (ThermoFisher Catalog # 21565) for 30 minutes. 4X Laemmli sample buffer supplemented with 10% beta-mercaptoethanol was added and sample were boiled briefly for 5 minutes to quench reactions. Samples were then resolved on a 6% SDS-PAGE gel and Western blot analysis for HSF-1 was carried out as described above.

### Whole Worm/Gonad Dissections Immunofluorescent staining

Anti-serotonin and anti-HSF1 staining was performed following the protocol developed by the Loer Lab (http://home.sandiego.edu/~cloer/loerlab/anti5htshort.html) and modifications were described in full in Tatum *et al*, 2015 (7). Briefly, worms were picked into 500ul of 1X PBS (pH 7.4) and spun down quickly then fixed in 4% paraformaldehyde (Catalog no. 15710; Electron Microscopy Sciences, Hatfield, PA) in 1X PBS (pH 7.4) at 4°C for 18 hours. Worms were then incubated in beta-mercaptoethanol solution for 18 hours, followed by cuticle digestion using collagenase type IV (Catalog no. C5138; Sigma, St. Louis, MO). For serotonin staining, the primary antibody was 1:100 rabbit anti-serotonin as in Tatum et al., 2015. Primary antibody used was 1:100 rabbit anti-HSF1 antibody (Sigma) while the secondary antibody used was 1:100 donkey anti-rabbit Alexa647 (Life Technologies Catalog No: A-31573). For staining of nuclear pore complex proteins, we used 1:100 mouse anti-NPC (54) (Abcam, ab24609) and 1:100 rabbit anti-mouse Alexa488 (Life Technologies Catalog No. A-11059). Worms were incubated with DAPI in 0.1% BSA/1X PBS for 30 minutes at room temperature then washed and mounted on slides in approximately 12 μl of Vectashield mounting medium (Catalog # H-1000; Vector Laboratories, Burlingame, CA). Imaging of slides was performed using a Leica TCS SPE Confocal Microscope (Leica, Solms, Germany) using a 63x oil objective. LAS AF software (Leica, Solms, Germany) was used to obtain and view z-stacks. For gonad dissections to stain for RNA Pol II, day one adult AM1061 worms were picked into 15ul of 1X PBS (pH 7.4) on a cover slip, and quickly dissected with a blade (Integra Miltex Product # 4-311, York, PA). A charged slide (Superfrost Plus, Fisher Scientific Catalog # 12-550-15, Pittsburgh, PA) was then placed over the cover slip and immediately placed on a pre-chilled freezing block on dry ice for at least 5 minutes. The cover slip was quickly removed and the slides were fixed in 100% methanol (-20°C) for 1 minute then fixed in 4% paraformaldehyde, 1X PBS (pH 7.4), 80mM HEPES (pH 7.4), 1.6mM MgSO4 and 0.8mM EDTA for 30 minutes. After rinsing in 1X PBST, slides were blocked for one hour in 1X PBST with 1% BSA and then incubated overnight in 1:500 mouse anti-Pol II (Biolegend, # MMS-126R clone 8w16g(76)). The next day, slides were washed and then incubated for 2 hours in 1:1000 donkey anti-mouse Cy3 (Jackson ImmunoResearch Lanoratories, Code # 715-165-150, West Grove, PA) before being washed and incubated in DAPI in 1X PBST, then mounted in 8ul Vectashield and imaged as described above. For quantification of nuclear bodies, the images were merged when co-staining was carried out (HSF-1 and NPC or HSF-1::GFP and Pol II) and discrete puncta ranging in size from approximately 400-550 nm were counted first in the HSF-1 channel. We then determined whether these HSF-1 puncta co-localized with the puncta in the other channel. When HSF-1 immunostaining alone was carried out, only the HSF-1 channel was used to quantify nuclear bodies.

### Optogenetic activation of serotonergic neurons (olfaction + choice assay)

To make experimental all*-trans* retinal (+ATR) plates, a 100mM ATR (product no. R2500; Sigma, St. Louis, MO) stock dissolved in 100% EtOH was diluted to a final concentration of 0.4 mM into OP50 and 250μl was seeded onto a fresh NGM plate. Control (-ATR) plates for experiments were seeded at the same time with the same OP50 culture, but without ATR. Plates were kept in dark and allowed to dry for a minimum of 10 hours prior to use. Plates were never used later than 1 day after they were seeded with ATR. The *C. elegans* strain used in this experiment was AQ2050 (*ijIs102; lite-1(ce314)*). L4s were harvested onto +ATR and –ATR plates, and the experiment was carried out on day one adult worms that were transferred in sets of 10 worms per plate onto plates containing 5ul of either +ATR or –ATR OP50 lawns. All plates were kept in the dark and animals were allowed to acclimatize to room temperature (20-22°C) for at least 30 minutes prior to the start of the optogenetic activation.

For the olfaction, HT115 and PA14 bacterial cultures were grown to an OD600 range of 1.4-1.7, and five 10ul drops of culture (either HT115 or PA14) were placed around the small bacterial lawn. The animals were then immediately illuminated with blue light for 5 minutes at a 6.3x magnification using a MZ10 F microscope (Leica, Solms, Germany) connected to a EL6000 light source (Leica, Solms, Germany), and subsequently transferred onto choice plates containing HT115 and PA14 lawns. –ATR animals were treated similarly. During the process of olfaction and optogenetic activation, animals that moved away from the central lawn were not used for the subsequent choice assay.

### Scoring germline nuclei for HSF-1::GFP activation post-olfaction

Olfaction was carried out as described above (“Olfactory pre-exposure/training”), using water and 2AA. The *C. elegans* strain used was AM1061 *(rmSi1 II; hsf-1(ok600) I)*. Whole worm live imaging was carried out using a Zeiss Observer A1 inverted microscope and animals were immobilized in 25mM levamisole on 2% agarose pads with cover slips. Nuclei were scored for induction within 10 minutes of the end of olfaction. Induction was assessed based on the presence or absence of HSF-1::GFP stress-induced nuclear puncta in the nuclei of germ-cells located in the two gonads of *C. elegans*. Analysis of the HSF1::GFP puncta was carried out by counting the number of nuclei showing distinct puncta compared to the total number of nuclei present in a single focal plane. Images also included animals put on OP50 (control) or PA14 for 30 minutes on PA14 at 20°C.

### Statistical Analysis and N values

For qRT-PCR data expressed as fold change in mRNA levels relative to control, a linear mixed model analysis for a randomized block design was used to compare the different conditions in each experimental data set. This was done to account for variation between different biological replicates, where treatment response was compared within the experiment. The data for this analysis was the response measure expressed as a ratio of control (for example, OP50 or H2O odor only). Since the distribution of ratios was usually not normally distributed, the natural log transformation was applied to the data to normalize the data distribution, with the log transformed values used in the analysis. Means in the log scale were then back-transformed to obtain geometric mean estimates in the original scale. The statistical analysis was performed using MIXED procedure in SAS (version 9.4). For all other data sets and where described (choice assays, smFISH/immunofluorescence/fluorescence quantification, Western blot quantification, ChIP-qPCR analysis), the parametric Student’s t-test (paired) was conducted to test for significance. For longevity/survival assays, OASIS software available online (https://sbi.postech.ac.kr/oasis2/surv/) was used to calculate mean lifespan, and the log-rank test was used to calculate statistical significance between the different conditions.

For all data quantified and analyzed, N values represent at least 3 independent repeats conducted to generate the number in each of the represented data points. For data that do not require quantification, the experiments were also conducted 3 or >3 times, and the data shown are representative data. N values for choice assays are the numbers of independent repeats of the Choice Assay. N values for qRT-PCR data are the number of repeats of the mRNA measurements per sample. When assays are conducted to compare the difference between different genotypes, only the number of paired repeats are considered to generate the N value. The N value in smFISH data, or data regarding HSF-1 nuclear bodies, is numbers of nuclei counted per animal (/field of observation) per genotype per treatment condition obtained from 2-3 independent repeats of the experiment. Only N of paired experiments are used when genotypes are compared.

### Chromatin Immunoprecipitation (ChIP)

Preparation of samples for ChIP were performed by modifying the protocols previously described (50, 77). 100 wild type day one adult animals per condition were collected from NGM plates, washed with 1X PBS (pH 7.4) and cross-linked with 2% formaldehyde at room temperature for 10 minutes. Reactions were quenched with 250mM Tris (pH 7.4) at room temperature for 10 minutes, and then washed 3 times in 1X PBS with protease inhibitor cocktail and snap-frozen in liquid nitrogen. The worm pellet was resuspended in FA buffer (50mM HEPES pH 7.4, 150mM NaCl, 1mM EDTA, 1% Triton X-100, 0.1% sodium deoxycholate) supplemented with protease inhibitor cocktail and lysed using a Precellys24 homogenizer (Bertin Corp., Rockville, MD), then sonicated in a Vibra-Cell processor (Sonics & Materials Inc., Newtown, CT). Pre-cleared lysate was then incubated overnight with 5ul of Rabbit anti-HSF-1 antibody (Sigma) and immunoprecipitation was performed with Protein A/G magnetic beads (Pierce, Catalog # 88802). qPCR analysis of DNA was performed using the reagents described above and the primer sets *syp-1* and *hsp-70* (F44E5.4) were used to respectively quantify non-specific and specific binding of gene promoters to HSF-1. The amplified qPCR products were run on agarose gels to verify that ChIP had resulted in the amplification of the appropriate sized band. PCR amplification of F44E5.4 was also carried out to visualize amplification of the appropriate product in an agarose gel.

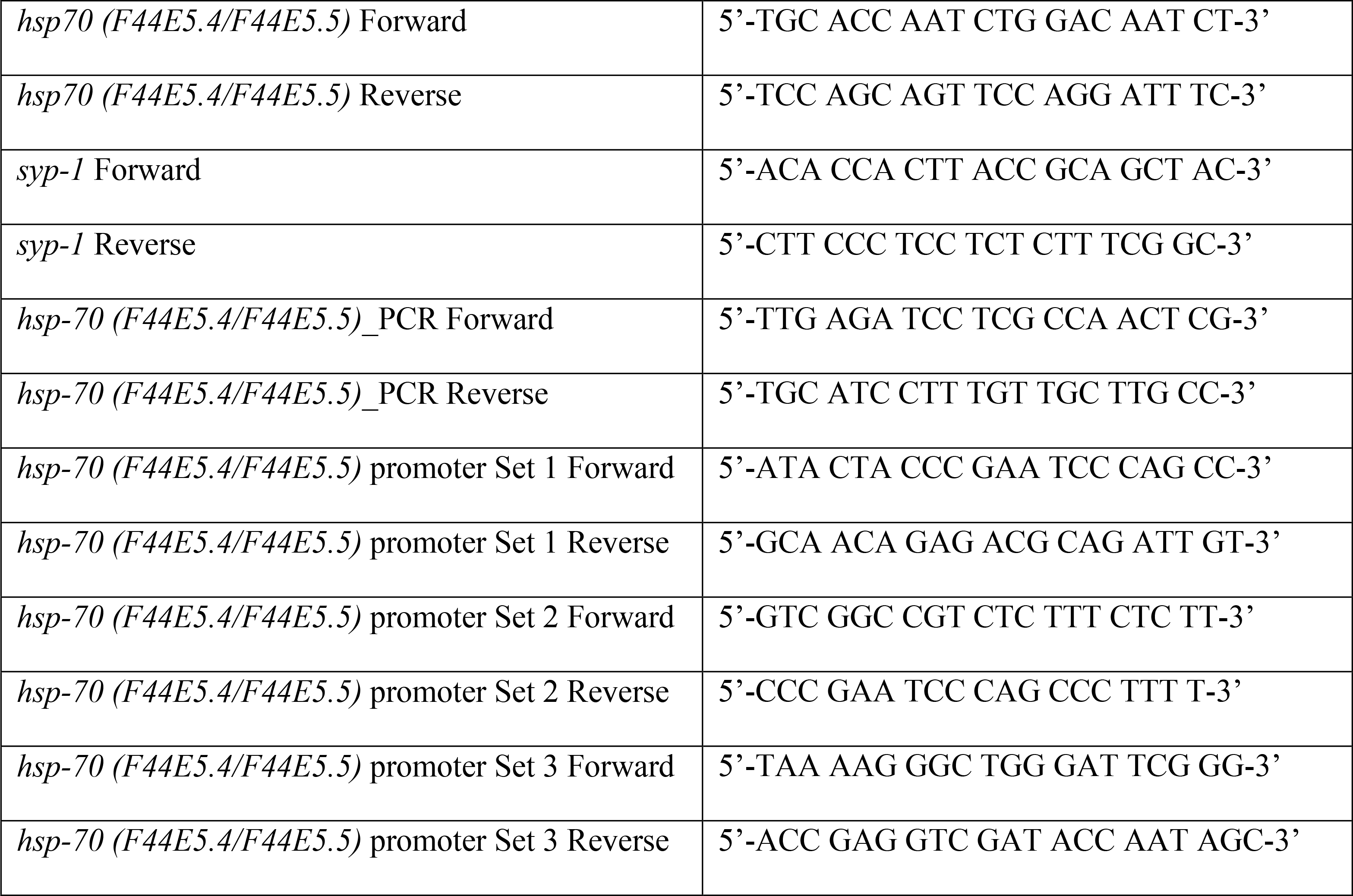

### Electrophoretic Mobility Shift Assay (EMSA)

HSE probe sequences were obtained from Catarina-Silva *et al*, 2013(78) and IR700-labeled oligos were obtained from Integrated DNA Technologies (Coralville, IA). Worm lysate was prepared by washing worms off in 1x PBS (pH 7.4) and immediately snap-freezing in liquid nitrogen. Worm pellets were thawed on ice and lysed in a binding buffer (10mM HEPES pH 7.4, 130mM NaCl, 5mM KCl, 1mM EDTA, 0.2% NP-40, 10% glycerol) supplemented with protease inhibitor cocktail and 1mM DTT. Lysis was carried out using the Precellys24 homogenizer (Bertin Corp., Rockville, MD). EMSA binding reactions (lysate, poly dI-dC and labeled IR700-HSE probes) were incubated at room temperature for 30 minutes, except for “heat shock” binding reactions, with or without competition using unlabeled probes, which were performed at 35°C for 30 minutes. Samples were then run out on a 6% acrylamide gel in 0.5X TBE and imaged using Li-Cor^®^ Odyssey Infrared Imaging System (LI-COR Biotechnology, Lincoln, NE) and quantified using Li-Cor^®^ Image Studio software.

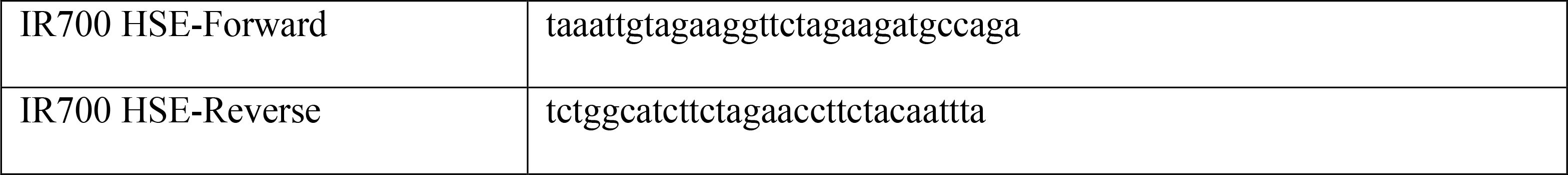

### Motility Assays

For motility assays, 2^nd^ generation RNAi animals were used (refer to growth conditions described above). Animals were harvested as L4s the day prior to the experiment. Day one adults were singled onto a lawn of OP50 and a video of the animals’ movement was captured at 0.8x magnification using a Leica MZ120 camera attached to an upright microscope (Leica KL1500) for approximately 30 seconds. Videos were analyzed using ImageJ software to measure the distance travelled by the animal and from this the velocity of the worm was calculated. The velocities were then used to calculate the time needed for each animal to travel 1 inch (the distance between bacterial lawns in the choice assay described above).

## Supplementary Materials

Fig. S1.

Design and specificity of olfactory pre-exposure and choice assay.

Fig. S2.

The compound 2-aminoacetophenone (2AA) made by PA14 specifically modulates olfactory avoidance behavior and protects against PA14-induced death.

Fig. S3.

Serotonin is required for learning-mediated HSF-1 activation.

Fig. S4.

Characterization of *C. elegans* HSF-1.

Fig. S5.

Characterization of *C. elegans* HSF-1 following exposure to water ‘odor’ and 2AA odor.

Fig. S6.

The formation of HSF-1 nuclear bodies does not require RNA Polymerase II.

Fig. S7.

HSF-1 is required for olfactory learning.

Table S1.

Survival of animals on PA14 is dependent on HSF-1.

Table S2.

Statistical analysis.

Table S3.

2AA does not appear to be toxic to *C. elegans*

Table S4.

Pre-exposure to the odor of PA14 protects animals from subsequent exposure to PA14.

## Acknowledgments

We would like to thank past and current members of the V.P. laboratory for comments and discussion, and in particular Matthew Wheat, Kat Dvorak and Charu Anbalagan for assistance with data generation. We would like to acknowledge Dr. Smolikove (University of Iowa) for sharing protocols and advice, Dr. Yahr (University of Iowa) and Dr. Aballay (Duke University) for PA14 strains, Dr. Ruvinsky (Northwestern University, Chicago) for helpful suggestions and advice, *Caenorhabditis* Genetics Center (funded by NIH Office of Research Infrastructure Programs (P40 OD010440) for worm and bacterial strains. **Funding:** F.K.O was supported by V.P’s grants, a graduate student fellowship from the Developmental Studies Hybridoma Bank (DSHB) and a scholarship from Glenn/AFAR Foundation. V.P. is funded by the National Institutes of Health (NIH R01 AG 050653) and The Lawrence Ellison Medical Foundation (AG-NS-1056-13). **Author contributions:** F.K.O. and V.P. designed the project and experiments, analyzed data and wrote the manuscript, F.K.O performed experiments. **Competing interests:** The authors declare that they have no competing interests. **Data and materials availability:** All worm strains, material and protocols will be made available on request

